# Insights into the diversity and survival strategies of soil bacterial isolates from the Atacama Desert

**DOI:** 10.1101/2020.09.24.312199

**Authors:** Alicyn Reverdy, Daniel Hathaway, Jessica Jha, Gabriel Michaels, Jeffrey Sullivan, Daniela Diaz Mac-Adoo, Carlos Riquelme, Yunrong Chai, Veronica G. Godoy

## Abstract

The Atacama Desert, the driest and oldest desert in the world, is a hostile environment for life. Despite the inhospitable conditions, bacterial sequences detected in this location suggest rich bacterial life. This study tested the idea that certain bacteria would thrive in this location and that some of them could be cultivated permitting further characterization. Environmental surface soil samples from 1-5 cm deep were collected from 18 diverse locations within the Atacama Desert. To assess the bacterial taxa, diversity, and abundance, Illumina 16S rRNA gene sequencing was performed directly on soil samples. Bacteria were also cultured from the samples. We have a collection of 74 unique bacterial isolates after cultivation and confirmation by 16S rRNA gene sequencing. Pigmentation, biofilm formation, antibiotic production against *Escherichia coli* MG1655 and *Staphylococcus aureus* HG003, and antibiotic resistance were assessed on these isolates. We found that approximately a third of the colonies produced pigments, 80% of isolates formed biofilms, many isolates had antibiotic activity against *E. coli* and/or *S. aureus,* and many were resistant to commercial antibiotics. The functional characterization of these isolates gives us insight into the adaptive bacterial strategies in harsh environments and enables us to learn about their possible use in agriculture, healthcare, or biotechnology.

**Originality-Significant Statement:** This study provides the first microbial diversity analysis from Atacama Desert soil, presents the cultivation and isolation of 74 unique bacterial isolates, many of which may be novel genera and species, and explores pigment production, antibiotic production and resistance, and unique biofilm development as bacterial survival strategies for living within extreme environments.

## INTRODUCTION

Extreme environments are challenging for life, yet bacteria are resourceful and have developed multiple strategies for survival. One location in the world characterized by its extreme environment is the Atacama Desert, located in the north of Chile at the border of Bolivia and Argentina spanning about 128,000 km^2^ (Rundel P 2007). Bound by mountain ranges that prevent precipitation, the Atacama Desert is the oldest and driest, temperate desert in the world (Bull et al. 2018; Azua-Bustos, Urrejola, and Vicuna 2012; Azua-Bustos, Caro-Lara, and Vicuna 2015). Described as an extremobiosphere, the hyper-arid Atacama Desert contains a chain of Andean volcanoes, large salt flats, high mineral deposits, and altiplano lakes (Bull et al. 2018; Orellana et al. 2018). Ranging between 2,000 and over 5,000 meters above sea level, the region experiences the highest levels of UV radiation in the world and low levels of oxygen, as well as large temperature fluctuations and extremely low precipitation levels (Orellana et al. 2018) (Bull et al. 2016) (Cordero et al. 2018; Azua-Bustos, Urrejola, and Vicuna 2012; Azua-Bustos and Gonzalez-Silva 2014; Demergasso 2010). Thus, the Atacama Desert is of increasing interest for understanding the microbial diversity and strategies to survive in extreme conditions including alkaline or acidic pH, temperature variabilities, water stress, and high UV radiation (Orellana et al. 2018), conditions that resemble those of other planets (Parro et al. 2011).

Extremophiles are bacteria that adjust to and survive in hostile environments once thought too harsh to sustain life (Rampelotto 2013). These bacteria are categorized into two types: those that require the extreme condition, and those that tolerate extreme conditions, but can also grow optimally in “standard” conditions (Rampelotto 2013). To thrive in harsh conditions, extremophiles use a variety of strategies to maintain their communities. These strategies have been extensively reviewed and include accumulation of certain molecules to counteract imbalances produced by the extreme conditions, production of specially designed enzymes and cell membranes, and enhanced DNA-repair mechanisms (Orellana et al. 2018) (Bull et al. 2016) (Azua-Bustos and Gonzalez-Silva 2014; Azua-Bustos, Urrejola, and Vicuna 2012).

Other such strategies include biofilm formation and antibiotic and pigment production (Hall-Stoodley, Costerton, and Stoodley 2004) (Azua-Bustos, Urrejola, and Vicuna 2012; Gerardin, Springer, and Kishony 2016). Biofilm is a community of surface attached bacterial cells encased in a self-produced, protective matrix that primarily consists of protein, exopolysaccharide, and extracellular DNA (Vlamakis et al. 2013; Hall-Stoodley, Costerton, and Stoodley 2004). Biofilms enhance nutrient sharing, cell-cell communication (Hall-Stoodley, Costerton, and Stoodley 2004; Vlamakis et al. 2013), and provide a protective barrier against antimicrobials, UV damage, and other pathogens (Hall-Stoodley, Costerton, and Stoodley 2004; Hall and Mah 2017; de Carvalho 2017). As such, biofilm-producing bacteria are one of the leading causes of human and plant pathogenesis, consistent with its role as a bacterial survival strategy (Vlamakis et al. 2013; Hall-Stoodley, Costerton, and Stoodley 2004).

Bacterial antibiotic production is a competitive advantage since it allows antibiotic producing bacteria to fend off other colonizers and protect the population from further harm (Gerardin, Springer, and Kishony 2016). For this reason, the search for novel antibiotics from environmental isolates is a very active area of research. The hope is to discover antibiotics to use against specific pathogenic bacteria without harming the general microbiome (Ferri et al. 2017) (Azua-Bustos and Gonzalez-Silva 2014) (Lewis 2017). Many groups have isolated and identified new compounds produced by *Actinobacteria* from the Atacama Desert, supporting the study of extremophiles for novel antibiotic discovery (Rateb, Houssen, Harrison, et al. 2011; Rateb, Houssen, Arnold, et al. 2011; Undabarrena et al. 2016; Goodfellow et al. 2018; Santhanam, Okoro, Rong, Huang, Bull, Andrews, et al. 2012; Santhanam, Okoro, Rong, Huang, Bull, Weon, et al. 2012; Okoro et al. 2009). But because bacteria are exposed to antimicrobials produced in their environment, they are apt to develop strategies for surviving these challenges (Ferri et al. 2017; Lewis 2017). Antibiotic resistance has created a billion-dollar problem in the healthcare system (Ferri et al. 2017). Moreover, identifying naturally resistant bacteria will provide us with mechanistic models to better understand how bacteria become antibiotic resistant as well as insights into how to combat this rapidly increasing problem.

Microbial diversity studies from the Atacama Desert have only been published within the last 15 years (Bull et al. 2016). Those that have focused on bacteria, have analyzed individual phyla such as *Cyanobacteria* and *Actinobacteria* (Bull et al. 2016). More recent publications have found the dominant phyla in the Atacama Desert to consist of *Actinobacteria, Proteobacteria,* and *Bacteriodetes* (Mandakovic, Maldonado, et al. 2018; Mandakovic, Rojas, et al. 2018; Dorador et al. 2018; Demergasso et al. 2004; Demergasso et al. 2008; Drees et al. 2006; Schulze-Makuch et al. 2018; Dorador et al. 2010; Demergasso 2010). Depending on the sampling location, sample type (water vs. soil), sample depth, and time of year, these communities change; though the overall function within a specific ecological landscape remains (Demergasso et al. 2004; Demergasso et al. 2008; Dorador et al. 2018; Mandakovic, Rojas, et al. 2018; Uritskiy and DiRuggiero 2019). Overall, there is still a large gap in knowledge of the microbial diversity in the Atacama Desert. With better sequencing technology and increased characterization of cultivable isolates, we will gain a more complete view of what and how extremophiles live in their environment. Additionally, investigating which bacterial communities live in different locations is essential for understanding how they contribute to the environmental ecology and drive global dynamics.

Here, we provide a comprehensive picture of the microbial diversity and investigation into the survival strategies of bacteria living in the Atacama Desert. We sampled from 18 locations within the Atacama Desert. By applying Illumina 16S rRNA gene sequencing directly on soil samples, we provide a picture of microbial diversity in this extreme environment. Further, we cultivated 74 unique isolates and performed characterization assays to identify pigment production, biofilm formation, antibiotic production, and antibiotic resistance as probable survival mechanisms. In this study we provide insights into bacterial diversity and the strategies that bacteria use to survive in extreme environments.

## RESULTS

### Locations and environmental sampling

The Atacama Desert exhibits the most extreme environment in Chile (Orellana et al. 2018; Bull et al. 2018). For this reason, we were curious to see what bacteria were present in order to identify what strategies they may use to survive in such a harsh environment. During the months of May and June of 2018, we travelled to 18 locations in the Atacama Desert (Figure 1, Figure S1). Each high-altitude location is distinct in its characteristics: fully barren to presence of low grasses and high to low mineral and salt content (Table S1, Figure S1).

**FIGURE 1.**
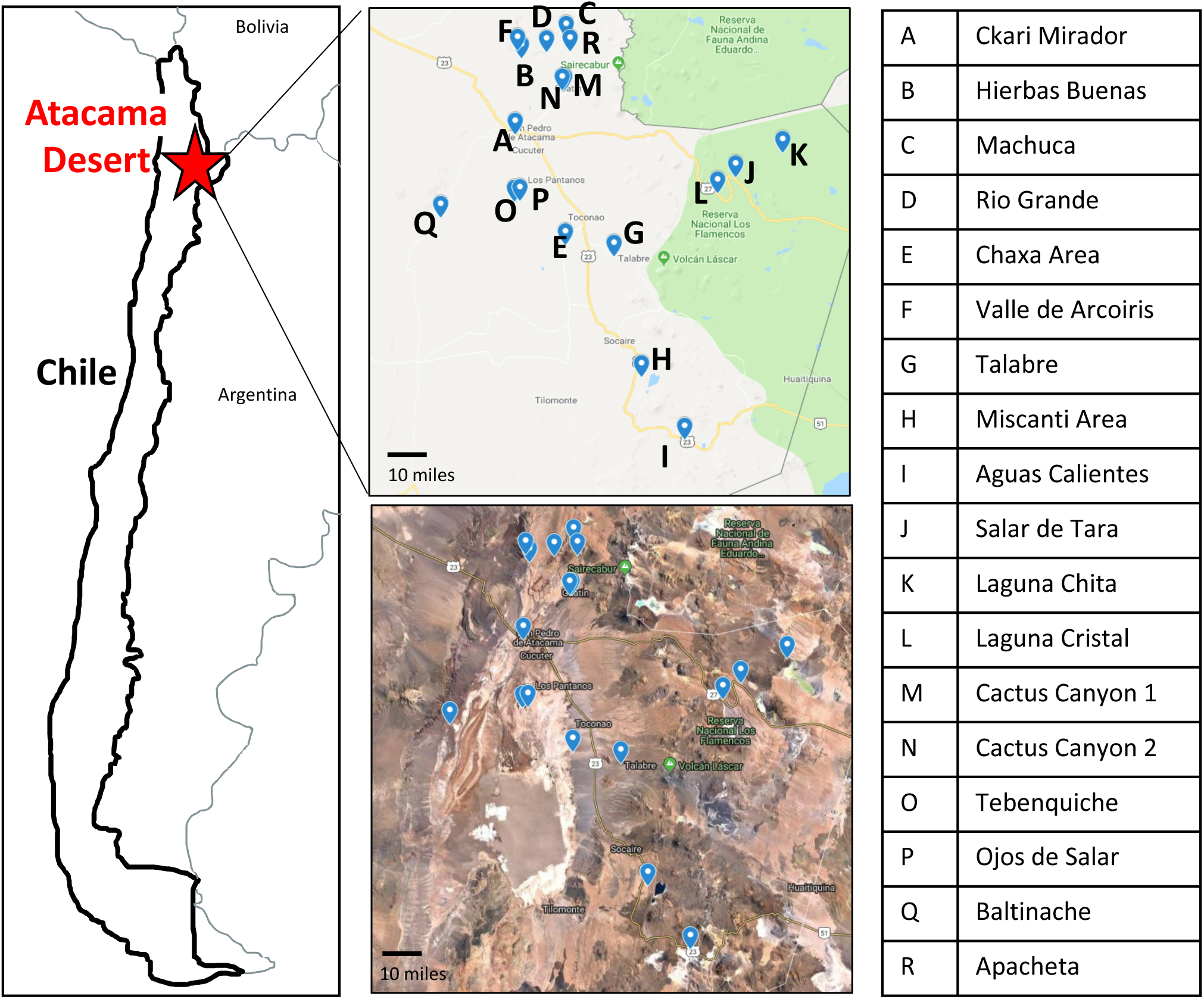
Map of the 18 sampling locations across the Atacama Desert. Sampling sites have clear geographical distinction as seen by Google Maps imaging. Letter key identifies the name of sampling location labeled on the maps.

The environment of the Atacama Desert is generally dry and barren (Figure 1). Indeed, the Atacama Desert is the oldest driest desert in the world and the least inhabited part of Chile (Bull et al. 2018). The average annual rain fall is 2 mm, which is extremely low when compared to other arid deserts (Azua-Bustos and Gonzalez-Silva 2014). Remarkably, air humidity is also kept low, given its geographical location bounded by two large mountain ranges, the Cordillera de los Andes to the east and the Coastal Cordillera to the west (Bull et al. 2018). Ranging between 2,000 and 5,000 meters above sea level, the Atacama Desert is also subjected to extreme levels of UV radiation and low oxygen content (Cordero et al. 2018; Escudero L 2007; Demergasso 2010). Large temperature fluctuations also occur, going from 0°C to 20°C in one day. During the year, the air temperature ranges from −6°C (21.2°F) to 38°C (100.4°F) (Azua-Bustos, Urrejola, and Vicuna 2012; Bull et al. 2018). The Atacama Desert contains many salt flats, which contributes to a high salt content and a high density of minerals including nitrates, iodates, boron, lithium, and copper deposits (Tapia et al. 2018). The Atacama Desert has been considered “a prime example of the extremobiosphere” (Bull et al. 2018). The ability for microbes to exist and survive under such extreme conditions makes them not only interesting to study, but key to understanding life at its limits.

Specific sampling locations were chosen for their unique characteristics; high altitude, near water, barren, high salinity, etc. At each location, environmental soil samples between 1-5 cm were collected. The exact coordinates were recorded at each sampling location (Table S1). Samples were then processed and analyzed to identify the total bacterial community content and to cultivate bacterial isolates (Figure 2).

**FIGURE 2.**
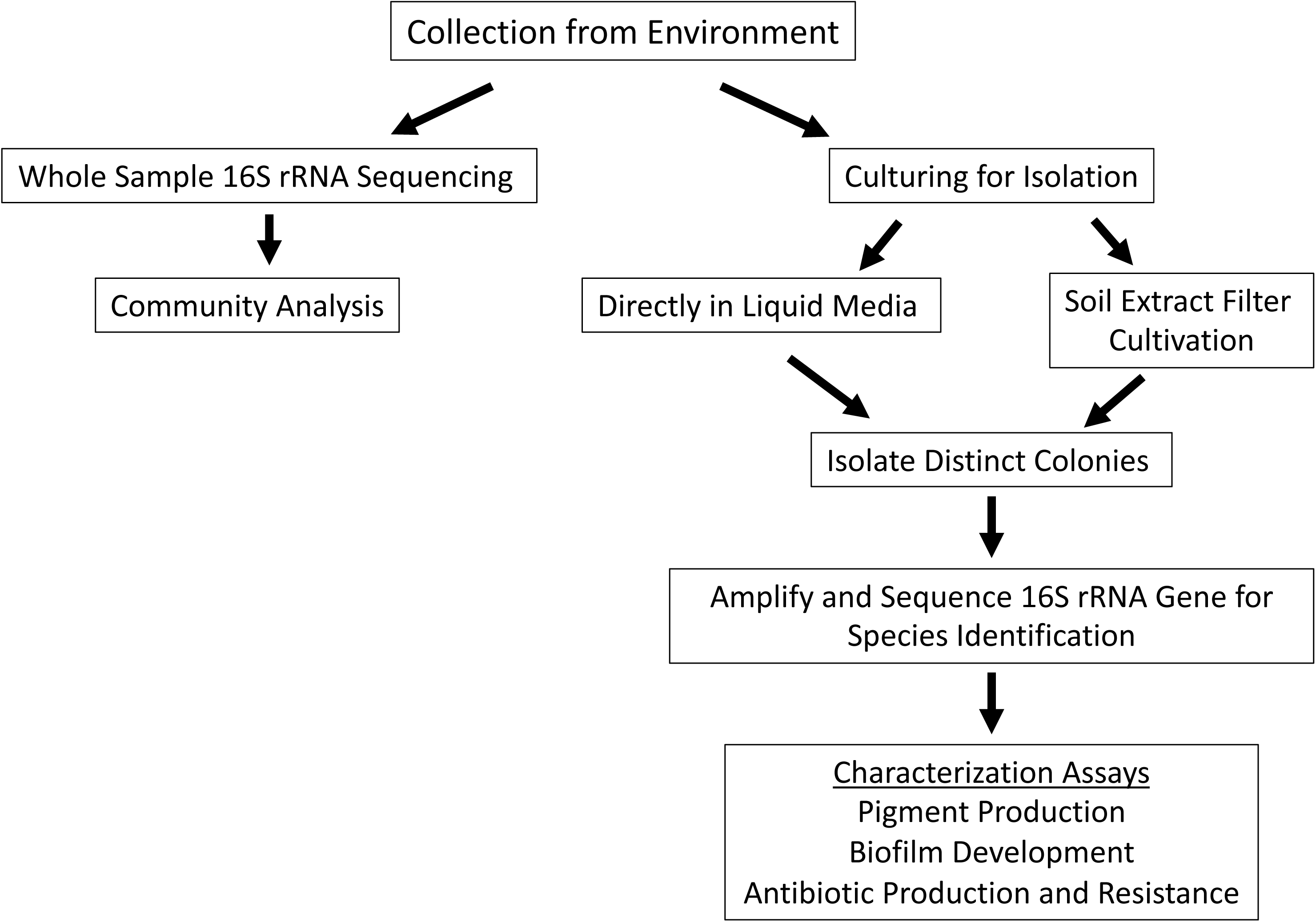
Schematic approach used for sample analysis and colony isolation. Samples were collected from locations in Figure 1. Illumina 16S rRNA sequencing was performed on soil samples to directly sequence and characterize the overall microbial ecology. Samples were also cultured to isolate individual bacterial species. This was done by two methods: inoculating the sample directly into culture medium or by scraping concentrated sample from soil extract filter disc and depositing it onto R2 agar plate. Bacterial colony growth was observed, and colonies were isolated until purity. Colony PCR of the 16S rRNA gene was used to identify species of the colony isolates with different morphologies. Colony isolates were then assayed for pigment production, biofilm production, and antibiotic production and resistance.

### Bacterial composition of the Atacama Desert

#### Archaea and Bacterial Abundance

To assess the microbial community composition of the samples, Illumina 16S rRNA sequencing was used directly on soil samples from the 18 sampling locations and analyzed for its Archaea and Bacteria composition. Importantly, the soil samples from Ojos de Salar and Baltinache (Figure S1P and Q) did not provide reliable data when compared to the negative control and could not be analyzed any further. This indicates that the soil composition was such that it did not permit the identification of microbial life with the test used. Community composition was successful in the remaining 16 soil samples. Of the total microbial content, it was found that 1.1% of the community was Archaea (Figure S2). This is comparable to levels found in the other hyper-arid desert soils (Bates et al. 2011; Schulze-Makuch et al. 2018).

#### Alpha Diversity

The focus of our search was on bacterial diversity. Looking at the number of different taxa in a sample, the alpha diversity, a measurement of the richness, ranged from 184 to 652 taxa (Figure 3A). Overall, this is low compared to other environments. Sea water has an estimated 10,000 species richness, the temperate forest and grasslands soil has a few thousand species, and the gut microbiome has about one thousand species (Kerrigan, Kirkpatrick, and D’Hondt 2019; George et al. 2019; Sears 2005; D’Argenio and Salvatore 2015). As predicted, our results suggest that the extreme environment has a significantly lower number of unique taxa detected using the direct sequencing method. Unfortunately, it is difficult to make direct comparisons with other studies as the sequencing methods have changed dramatically since these started.

**FIGURE 3.**
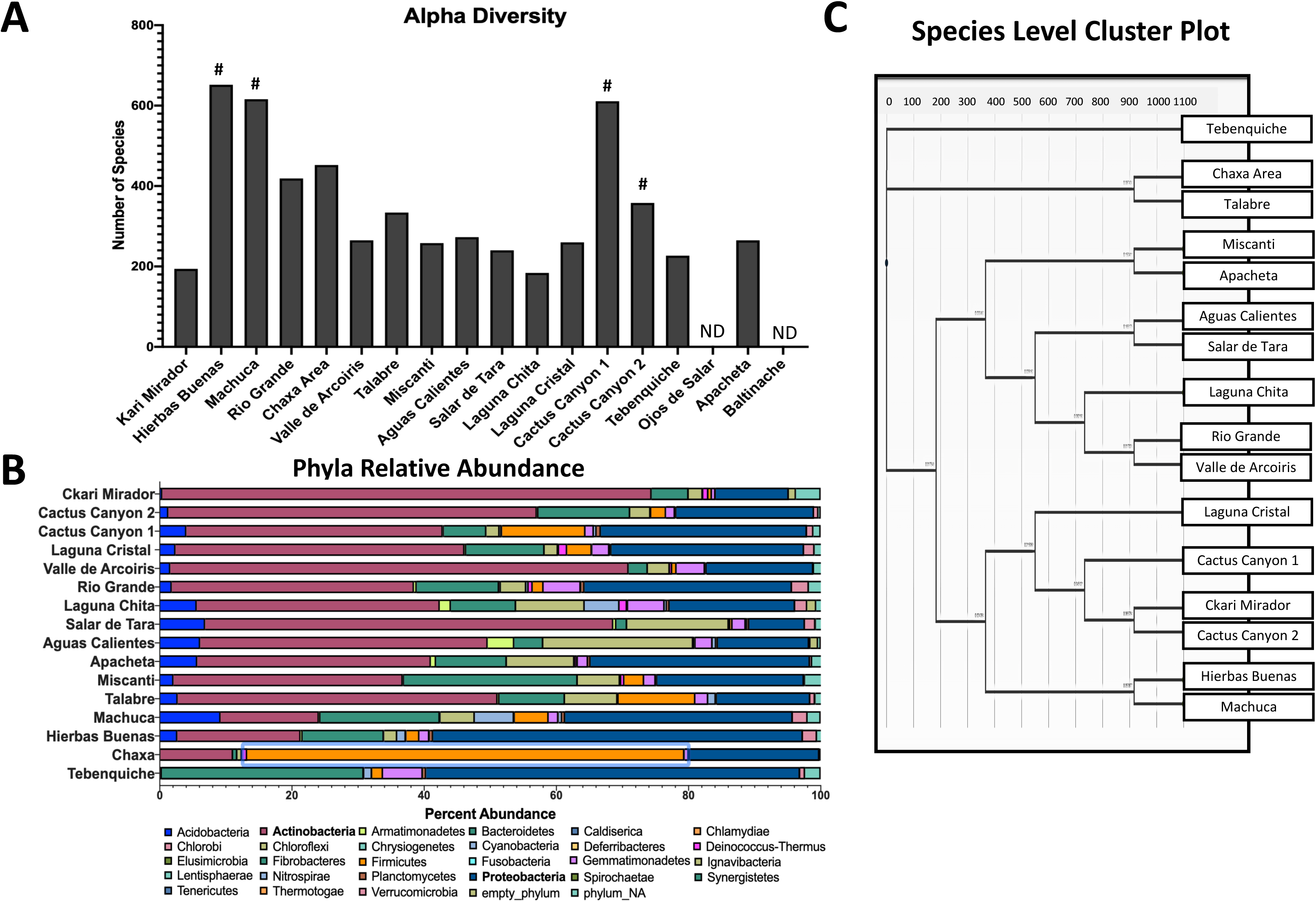
Illumina 16S rRNA sequencing provides a snapshot of bacterial diversity. (A) Quantification of alpha diversity soil samples. Each bar represents the number of unique taxa detected in each soil sample. Pound signs indicate locations of higher plant life. No detection of sequences in Ojos de Salar or Baltinache (ND). (B) Bar chart of the phyla relative abundance within each sample. Each color represents a different phylum and the length of the bar represents the percent abundance within that sample. *Actinobacteria* (pink) and *Proteobacteria* (dark blue) were the most abundant phyla. (C) Cluster plot dendrogram grouping samples based on genus similarities under Marisita-Horn analysis. Length of bar represents degree of similarity. Tebenquiche was the most different from the rest of the samples.

We find that locations that have more plant life, had a greater richness (Figure 3A). Hierbas Buenas (Good Herbs) has an abundance of low grass and bushes, and accordingly, had the largest richness at 652 taxa (Figure 3A, Figure S1B). Conversely, the locations with the driest and most barren soil had the lowest richness as seen at Laguna Chita, Ckari Mirador, Tebenquiche, Salar de Tara, and Valle de Arcoiris with 184, 194, 227, 240, and 265 taxa, respectively (Figure 3A, Figure S1R, A, O, J, and F). The correlation of “plant-rich” areas containing higher bacterial richness is expected due to the strong symbiosis and known importance that the bacterial rhizobiome has for plant growth and survival (S 2018).

#### Phyla Relative Abundance

The two dominant phyla across all locations were *Actinobacteria* at 35.9% (pink bars) and *Proteobacteria* (dark blue bars) at 26.7% (Figure 3B, Table S2A). Our results are consistent with other studies that have investigated the bacterial content in the Atacama Desert (Demergasso et al. 2008; Aguayo et al. 2017; Schulze-Makuch et al. 2018). Other studies have also found *Bacteroidetes* and *Cyanobacteria* to be the dominant phyla, however they were sampling water, while we sampled soil (Demergasso 2010; Mandakovic, Maldonado, et al. 2018; Dorador et al. 2018). *Bacteroidetes* is the third most abundant phylum across all the samples at 11.3% abundance (Figure 3B, Table S2A). At the genus level, the most abundant was *Arthrobacter* (total percent in all samples 5.7%), found in all samples except for Chaxa (Figure S1E), followed by *Bacillus* (total percent in all samples 4.3%) (Table S2B) found everywhere (Table S2B).

Looking at specific locations, the two salt flats, Chaxa and Tebenquiche (Figure S1E and O), have different abundance distributions compared to the other locations. Chaxa is within the Salar de Atacama and has very low bacterial diversity; 96.7% of its taxa were within *Actinobacteria, Proteobacteria*, and *Firmicutes*, and all other phyla contributed to less than 1% of the total bacteria (Table S2A). The high *Firmicutes* abundance is consistent with another study investigating the diversity in this location (Dorador et al. 2018). The *Firmicutes* phylum contains a high number of halophiles and spore-formers, and are known to have exceptional resistance to environmental stresses (Schulze-Makuch et al. 2018).

Tebenquiche, a large hyper-saline and high-altitude lake within the Salar de Atacama and known for its bacterial heterogeneity, has a unique phylum profile (Demergasso et al. 2004; Demergasso et al. 2008). Interestingly, this location contains the greatest abundance of *Bacteroidetes* at 30.6% compared to the average of 11.0% (Table S2A). This taxa distribution is consistent with analyses from other studies (Demergasso et al. 2004; Demergasso et al. 2008), (Oren 2013). Also of note, is that of the 227 unique species, there is a striking abundance of *Salinibacter* (21.4%) (Table S2B). *Salinibacter* is only detected at Tebenquiche (Table S2B). All other locations have a large species diversity, thus making this high singular species abundance and specific localization of interest.

#### Detected Isolates of Interest

It is important to note that 1.21% of the total bacterial content within the soil samples were unidentified phyla (Table S2A). Ckari Mirador (Figure S1A) had the highest percentage of unidentified phyla at 3.75% (Table S2A). This group of unidentified sequences may contain new species yet to be identified and characterized.

Other specific genera of interest, because they are poorly understood, are those that are found in high relative abundance (>10%) within their location (Table S2B). Genus *Segetibacter* was found at 21.5% in Miscanti. Genus *Marinicella* was found in high abundance at 15.0% in Talabre (Figure S1G). In Salar de Tara (Figure S1J), genus *Crossiella* and the unidentified family of order *Solirubrobacterales* were found in high abundance at 11.6% and 11.1%, respectively. Each genus has only been characterized in the last 15 years and only contains one to three identified species (Labeda 2001; Wang et al. 2018; Weon et al. 2010; Seki et al. 2015). Further, *Marinicella* has only been found in seawater and here, we detected it in soil from a location that is characteristically dry and kilometers away from the ocean. Further investigation of these bacteria will provide insights into the selectivity by the location and calls importance to the ecosystem dynamic.

#### Cluster Analysis

To compare the bacterial diversity across the different sampling locations, we generated a genus level cluster plot (Figure 3C). The salt flat, Tebenquiche, was the outlier group and the other salt flat, Chaxa, was in another outlier grouping with Talabre (Figure S1O, E, G). The clustering of Apacheta and Miscanti is most likely due to their high altitude, shrub-containing characteristics (Table S1, Figure S1R and H). Salar de Tara and Aguas Calientes are closely located and near high altitude lakes (Figure 1, Figure S1J and I)). Rio Grande and Valle de Arcoiris are two very dry and barren locations (Figure S1D and F). The clustering based on location similarities highlights that the genera are most likely selected by the environmental conditions.

#### Valle de Arcoiris Soil Analysis

Looking within a specific location’s bacterial diversity allows us to dive deeper into the understanding of what geophysical properties can support specific bacterial growth. Valle de Arcoiris (Rainbow Valley) is one such location that contains geophysical properties of interest (Figure S1F). Named for its different colored soils, it is an area enriched in clays, minerals, and metals (Figure 4A).

**FIGURE 4.**
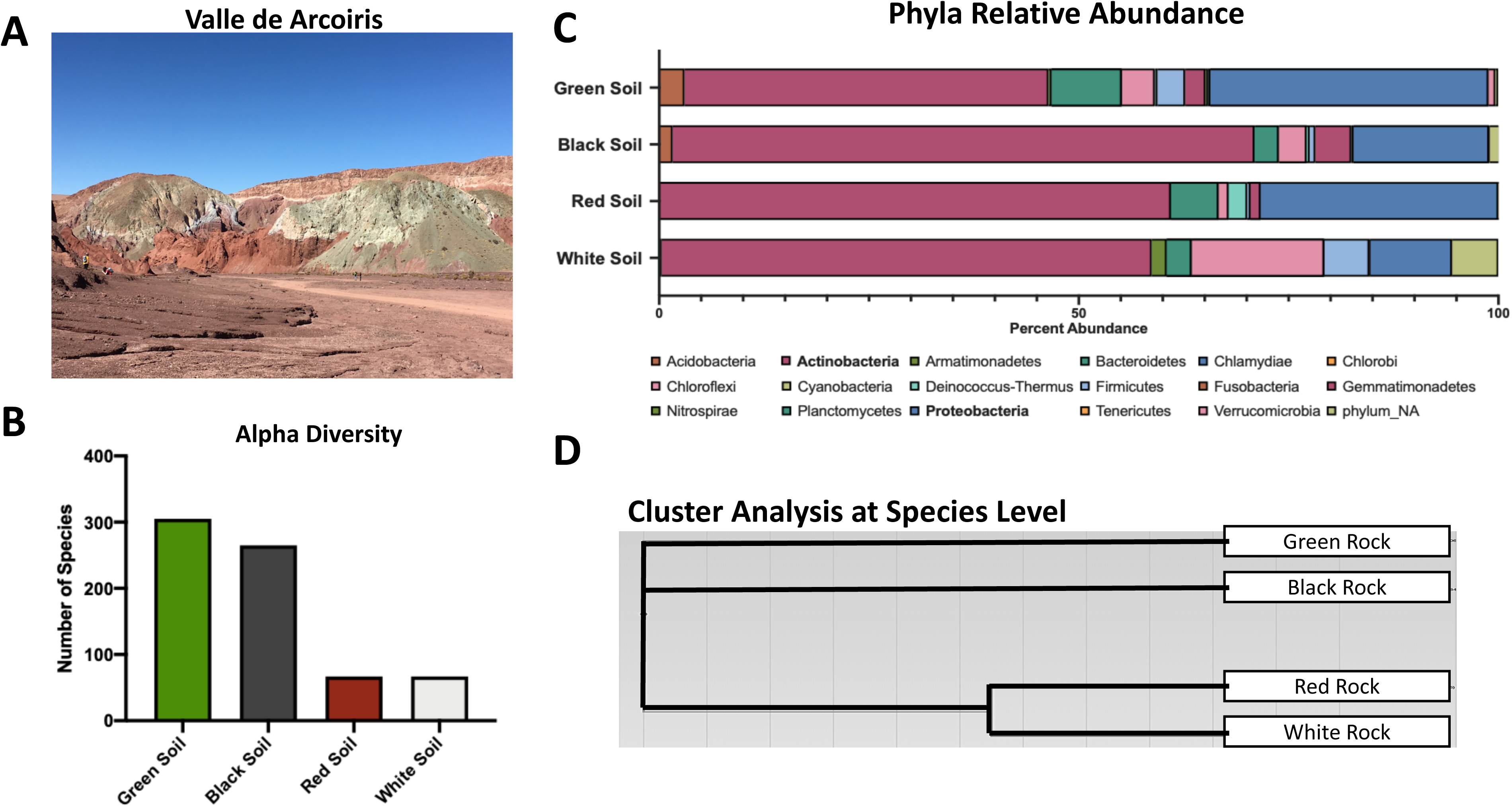
Analysis of Valle de Arcoiris soil samples demonstrate different bacterial diversity. (A) Image of the Valle de Arcoiris demonstrating the different colored soils. (B) Quantification of alpha diversity in each soil type. Each bar represents the number of unique taxa detected in each sample. (C) Bar chart of all the phyla relative abundance within each soil sample. Each color represents a different phylum and the length of the bar represents the percent abundance within that sample. (D) Cluster plot dendrogram grouping samples based on species similarities under Marisita-Horn analysis. Length of bar represents degree of similarity. The green and black soils and the red and white soils were demonstrated to be more similar, respectively.

The bacterial diversity was analyzed in 5 different soils; red, green, black, white, and white mineral. The two latter soils are of white color, but they are different since what we called “white mineral” soil did not amplify compared to negative control. This is indicative of undetectable microbial life in this soil sample. The others had sufficient amplification for comparative analysis. The green soil had the largest richness at 305 taxa, closely followed by the black soil at 265 taxa (Figure 4B). The red and white soils, however had a significantly reduced richness with both at 67 unique taxa. This indicates that either the mineral contents of the red and white soils are selective and preclude bacterial diversity, or that the detection method fails to find bacteria as the soil contents may be inhibitory to the methodology used.

Remarkably, there are no analyses that we could find indicating what specific salts were present in the different colored soils. It is likely that the soil color is given by the richness of transition metals such as Copper, Nickel, Cobalt, Iron, and Manganese in association with alkali metals such as Lithium, Sodium, and Potassium. All these salts are known to exist in the soils of the Atacama (Tapia et al. 2018). The green soil is believed to have an enrichment of magnesium, cobalt, and sulfur, while the red soil contains an enrichment of iron, and the white soil an enrichment of sodium chloride and calcium from shell erosion, among multiple other metals and minerals. More studies are needed in this area to advance the understanding of the Atacama soil microbiome.

As seen in the phyla relative abundance comparison, the percentage of *Proteobacteria* is different across soils in the Valle de Arcoiris (red 28.3%, black 16.1%, white 9.7%, green 33.1%) (Table S2C-1) indicating that this could be the phyla that is impacted the most by the differences in soil composition. Compared to the other three soils, the white soil had the most significant difference in abundance. Interestingly, the white soil had 5.5% of bacteria not identified to a known phylum, compared to the other three soils at <0.05%, 1%, and 0.3% (Table S2C-1). It is also the highest percentage of unknowns across all the Atacama Desert soil samples. This suggests several novel bacteria are contained in the white soil sample. Overall, the examples we find demonstrate an apparent pattern that at the genus level, what survives in the white soil is not found in the red soil, and vice versa (Figure 4D) (Table S2C-2).

Together, the microbial phyla and genera abundances provide insights in the taxa that are selected for by the components in these soils. These bacteria could also be the primary metabolizers that produce the different metals and minerals and thereby contribute to the specific soil composition. Further investigation of these bacteria will provide more insight into the dynamics between the bacteria and their ecosystem.

### Cultivation and isolation of unique colonies

Our next goal for gaining insights into the behavior of bacteria in the Atacama Desert was to cultivate and isolate single bacterial colonies from the samples that we collected. As indicated in detail in the materials and methods section, we used lysogeny broth (LB), diluted LB medium (1:100), and a low nutrient medium (R2) (Berdy et al. 2017). We found that, in general, isolates grew much better on the R2 medium than on the LB or 1:100 LB media plates. This was to be expected because bacteria from these locations are more accustomed to growing in low nutrient environments. Colonies were then purified on solid medium based on visual morphology differences. Two approaches were taken for cultivation (Figure 2). One approach involved arbitrarily adding ∼100 mg samples directly to cultivation medium either to liquid medium or spread on solid medium plates and incubated for a day at room temperature. The liquid-grown cultures were deposited on the surfaces of the same solid medium plates to allow for colony formation.

The other method for cultivation used a soil filtration method. Briefly, soil samples were resuspended in sterile water, filtered first through a filter paper to eliminate large size particles, and then through 0.22μm vacuum filtration system, and any trapped bacteria on the filter was kept for cultivation. To cultivate, a sterile stick scraped the surface of the filter disc, which was then resuspended into sterile water. The resuspension was then immediately spread onto R2 solid agar plates to allow for colony growth and further isolation. Colonies were grown at room temperature and were purified to a single colony morphology.

All the isolates from both cultivation methods were combined into a single set, and an initial screening among isolates from the same location was done to remove any isolates with the same colony morphology. This was performed to remove any potential sister colonies and resampling. For species identification, the 16S ribosomal gene was amplified and sequenced for all isolates.

It is important to note that we were not able to isolate any colonies from the Baltinache samples (Figure S1Q). This is consistent with the lack of amplification in the 16S metagenomic data set. This indicates that the Baltinache soil is either not hospitable to bacterial life, or our cultivation conditions were not conducive for the growth of bacteria that inhabit this soil. Baltinache is a salt flat, and the high salinity was not represented in our cultivation methods.

Results of the 16S rRNA gene sequencing identified 74 unique strains out of the 142 isolates (52%) across all sample locations (Table S3). Those strains belonged to 27 unique genera (19%) and 74 unique species (52%). Unique was defined as any isolate different to another isolate within a given sample. For example, if two isolates were identified to be *Enterobacter sp.* from the Cactus Canyon 1 location, but one was from a soil sample and another was from a moss sample, then they were considered unique. However, if two *Enterobacter sp.* isolates from Cactus Canyon 1 were both from a soil sample and both had a percent identification above 97% by 16SrRNA, then they were considered sisters and non-unique. A distribution of the unique and non-unique isolates from the different sampling locations was made using the above definition (Figure 5A).

**FIGURE 5.**
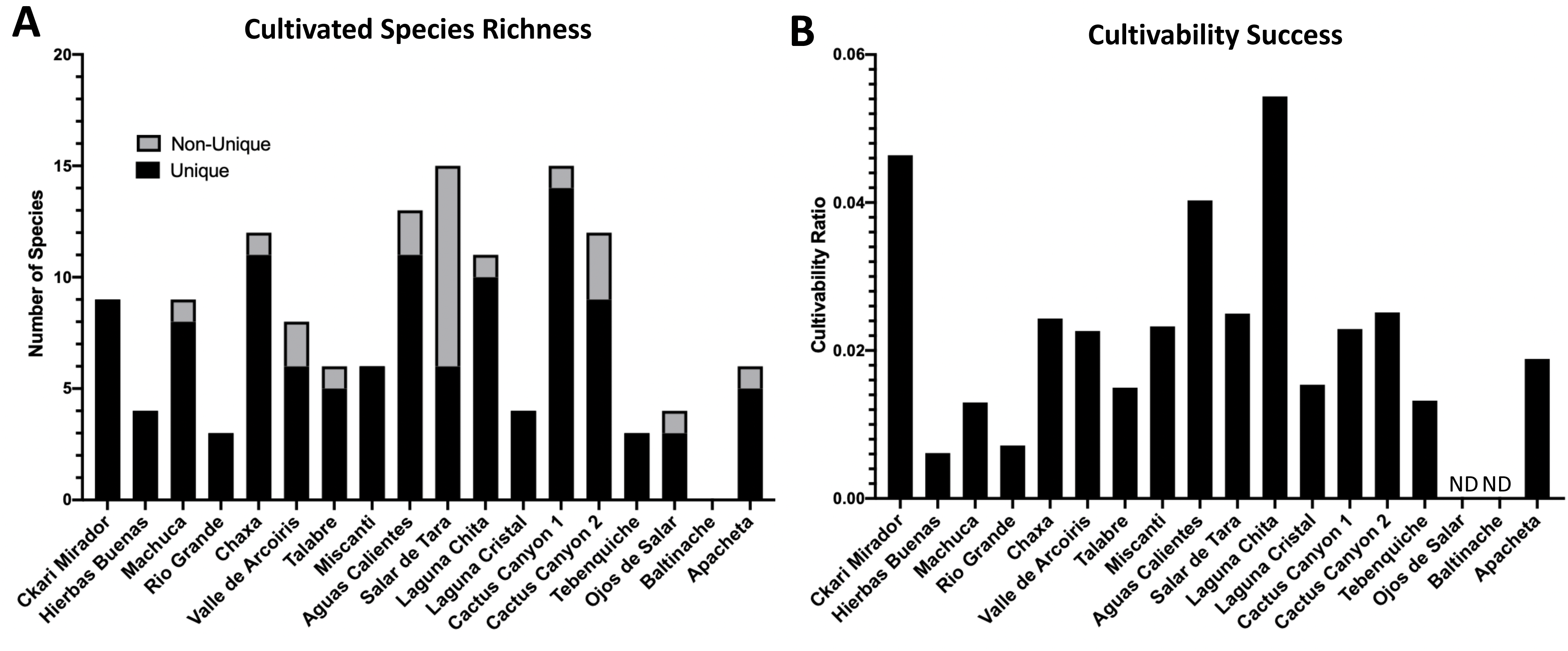
Among cultivable bacteria 74 unique colony species were isolated and identified. 142 distinct isolates based on colony morphology were isolated including 74 unique bacterial species. Isolates were cultured from environmental samples as described in Methods. The strain identity of colony isolates with different morphologies were distinguished based on the 16S rRNA gene of each colony isolate amplified via PCR and sequenced. The 16S rRNA gene sequence was aligned to the NCBI database to identify colony genus, species and/or strain using 97% identity as species separation and confidence. (A) Distribution of isolates across sampling locations. Unique is defined as a bacterial species within the same location that has no other representative. Non-unique refers to the same species found in different samples from the same location. (B) The cultivability success was calculated as the ratio between the cultivated species richness (unique) and the Illumina 16S rRNA alpha diversity (Figure 3A). Ojos de Salar and Baltinache could not have ratio calculated due to failure to detect bacteria in the Illumina data (labeled NA). Laguna Chita had the greatest cultivability success across all locations.

Based on Figure 5A, Cactus Canyon 1 had the greatest cultivability and diversity and Salar de Tara had the lowest diversity (Figure S1M, J). Chaxa, Aguas Calientes, and Cactus Canyon 1 had the best cultivability, and Rio Grande and Tebenquiche had the lowest cultivability based on our methods (Figure 5A, Figure S1E, I, M, D, O). To compare our cultivation success to the total amount of possible bacteria to cultivate from each soil sample, we calculated a cultivability ratio.

This is the number of unique cultivated species (Figure 5A) divided by the alpha diversity detected by the Illumina 16S rRNA sequencing directly from the full soil sample (Figure 3A). These data demonstrated that the bacteria from Laguna Chita, followed by Ckari Mirador and Aguas Calientes, had the greatest success in cultivation with our methods (Figure 5B, Figure S1K, A, I)). Consistent with the 16S rRNA sequencing, we could not generate a ratio for Baltinache (Figure S1Q) because there was no detection by sequencing, nor was there any cultivation. A ratio also could not be calculated for Ojos de Salar (Figure S1P) because the 16S rRNA sequencing did not accurately detect bacteria in the sample; however, we were able to cultivate three unique isolates from this location (Figure 5A). This finding suggests that there may have been inhibitory compounds in the soil sample from Ojos del Salar that did not allow for sufficient DNA amplification that is required by the methodology used.

For species identification, 97% was used as the species separation and confidence interval that a species was identified correctly, though 98.65% has also been used (Yoon et al. 2017). In our case, any percent identity below 97% indicates that the isolate is a novel species, which we may be undercounting. Interestingly, 34% of our isolates had a percent identity below 97%, and 4% of all isolates had a percent identity below 90%. This suggests that we may have at least 48 new species within our collection.

Of all the isolates, the most abundant genus was *Bacillus sp*. Forty-three isolates belonged to this genus and it was found in 13 locations (Table S3). *Bacillus simple*x was the most abundant species found in 10 locations (Table S3, Table 1). The next abundant genus was *Arthrobacter.* Fourteen isolates (9.8% of all isolates) were found in 7 locations (Table S3). These data demonstrate that *Bacillus* and *Arthrobacter* are non-discriminatory and can live in many diverse locations. These abundances also match the most abundant genera in the Illumina 16S sequencing data (Table S2B).

**TABLE 1.** Cultivated known plant-associated isolates.

| Species | ID (%) | Location | Reference |
| --- | --- | --- | --- |
| *Aeromonas aquatica* strain MX16A | 98.00 | Laguna Chita | (Beaz-Hidalgo et al. 2015) |
| *Agrobacterium rhizogenes* strain K599 | 89.57 | Valle de Arcoiris | (Valdes Franco et al. 2016) |
| *Aeromonas* sp. AU20 | 93.19 | Salar de Tara | (Anda et al. 2015) |
| *Bacillus atropheus* SQA-17 | 96.39, 99.05, 100 | Ckari Mirador, Cactus Canyon 2, Laguna Chita |  |
| *Bacillus atropheus* strain GQJK17 | 97.99, 98.03, 98.68 | Valle del Arcoiris, Laguna Chita, Chaxa, | (Ma et al. 2018) |
| *Bacillus megaterium* strain JX285 | 100 | Miscanti | (Huang et al. 2019) |
| *Bacillus mycoides* strain Gnyt1 | 99.71 | Miscanti | (Xiaomei Y 2020) |
| *Bacillus simplex* NBRC 15720 = DSM 1321 | 99.03, 98.19, 99.61, 99.13, 98.33, 88.70, 95.77<br>99.28, 94.67, 96.15, 98.61<br>98.49, 97.48, 99.01, 98.49,<br>98.31, 98.60, 95.69, 87.26,<br>94.37, 97.75, 99.81 | Talabre, Cactus Canyon, Salar de Tara, Cactus Canyon 1, Laguna Chita, Rio Grande, Valle del Arcoiris, Aguas Calientes, Ckari Mirador, Miscanti | (Gutiérrez-Luna 2010) |
| *Bacillus* sp. X1(2014) | 97.31 | Miscanti |  |
| *Bacillus subtilis* strain DKU_NT_03 | 99.99, 99.81, 95.90, 97.22, 96.50 | Talabre, Aguas Calientes, Salar de Tara, Cactus Canyon 1, Laguna Cristal | (Gutiérrez-Luna 2010) |
| *Bacillus subtilis* strain SG6 | 97.20 | Cactus Canyon 1 | (Zhao et al. 2014) |
| *Bacillus thuringiensis* strain c25 | 97.93 | Ojos de Salar | (Shrestha 2015) |
| *Bacillus thuringiensis* strain SCG04-02 | 97.93 | Ojos de Salar | (Fu et al. 2017) |
| *Brevundimonas* sp. LM2 | 96.79, 95.91 | Chaxa, Cactus Canyon 1 | (Chen et al. 2020) |
| *Enterobacter* sp. | 98.93, 93.82, 98.54, 98.62, 98.53, 98.66, 99.90, 99.90 | Cactus Canyon 1, Cactus Canyon 2, Tebenquiche, Laguna Cristal, Laguna Chita, Chaxa | (Taghavi et al. 2010) |
| *Janthinobacterium agaricidamnosum* NBRC102515 = DSM 9628 | 96.58 | Ckari Mirador | (Graupner, Lackner, and Hertweck 2015) |
| *Microbacterium oryzae* strain MB-10 | 99.90 | Cactus Canyon 2 | (Deb and Das 2020) |
| *Planococcus rifietoensis* strain M8 | 98.14, 97.01 | Cactus Canyon 2, Chaxa | (See-Too 2016) |
| *Pseudomonas azotoformans* strain S4 | 97.15, 99.26 | Tebenquiche, Rio Grande | (Fang et al. 2016) |
| *Pseudomonas brassicacearum* strain L13-6-12 | 99.70 | Hierbas Buenas | (Zachow et al. 2017) |
| *Pseudomonas fluorescens* NCIMB 11764 | 87.03 | Cactus Canyon 1 | (Silby et al. 2009) |
| *Pseudomonas koreensis* strain D26 | 98.32 | Tebenquiche | (Lin et al. 2016) |
| *Pseudomonas protegens* strain FDAARGOS 307 | 99.90 | Cactus Canyon 1 | (Ramette et al. 2011) |
| *Pseudomonas* sp. L10.10 | 97.40 | Laguna Chita | (See-Too et al.) |
| *Pseudomonas* sp. M30-35 | 96.10 | Laguna Chita | (He et al. 2018) |
| *Rahnella aquatilis* strain HX2 | 99.32 | Machuca | (Guo et al. 2012) |
| *Serratia fonticola* strain GS2 | 96.52 | Chaxa | (Jung et al. 2017) |
| *Serratia liquefaciens* strain FDAARGOS_125 | 99.80 | Laguna Chita | (Caneschi et al. 2019) |

From our list of identified isolates, we found 28 unique species that are known to be plant-associated (7 species contain isolates with identities < 97%) (Table 1, Table S3). They were found in all locations, except for Apacheta (Figure S1R). Some are beneficial and examples include *Bacillus subtilis, Pseudomonas brassicacearum* strain L13-6-12, *Serratia fonticola* strain GS2, and *Rahnella aquatilis* HX2 (Jeong et al. 2018; Vlamakis et al. 2013; Zachow et al. 2017; Jung et al. 2017; Guo et al. 2012). Some non-beneficial species include *Janthinobacterium agaricidamnosum* and *Pseudomonas koreensis* (Graupner, Lackner, and Hertweck 2015; Lin et al. 2016). Another species of note is *Enterobacter sp.* This is a plant-associated bacterium, but is also within the list of ESKAPE pathogens, which are nosocomial human pathogens that exhibit multi-drug resistance and can cause severe and lethal infections (Santajit and Indrawattana 2016; Taghavi et al. 2010). These groups of plant-associated bacteria may provide insight into how plants survive in these extreme environments.

Together, our data demonstrates success in bacterial cultivability from the extreme environments of the Atacama Desert. These isolates are representatives of where they came from and allow for the characterization of unknown species and identification of whether any of the known species have unique characteristics because they were isolated from an extreme environment.

### Pigment producing isolates

With the cultivated, isolated, and identified species from the Atacama Desert, we wanted to find out what these extremophiles could do. By identifying certain characteristics of these bacteria, we gain insights into their strategies for survival in these environments. The first easily identified characteristic was that 34.5% of our isolates produced a pigment (Figure 6A and 6B), most of which were cell bound. Of these isolates, 69.4% produced a yellow pigment, 26.5% produced an orange pigment, and 2 isolates produced a pink pigment. Pigment production by bacteria is thought to be a protective mechanism against UV radiation (Azua-Bustos, Urrejola, and Vicuna 2012; Narsing Rao, Xiao, and Li 2017) (de Carvalho 2017; Ordenes-Aenishanslins et al. 2016). Many *Actinobacteria* and *Cyanobacteria* from salt flats and hyper-arid locations produce protective pigments (Bull and Asenjo 2013; Narsing Rao, Xiao, and Li 2017). The Atacama Desert is known for its extreme levels of UV radiation (Demergasso 2010; Cordero et al. 2018). It is plausible that these bacteria produce pigments to protect themselves from the DNA-damaging UV that they are exposed to while living in the Atacama Desert. We attempted to measure the UV resistance of some of our isolates, but it is a difficult endeavor since the pigments are secondary metabolites produced late in stationary phase. In fact, most of the isolates tested were UV-sensitive when growing exponentially compared to *Escherichia coli* (data not shown). Pigment isolation would allow determination of its functional role.

**FIGURE 6.**
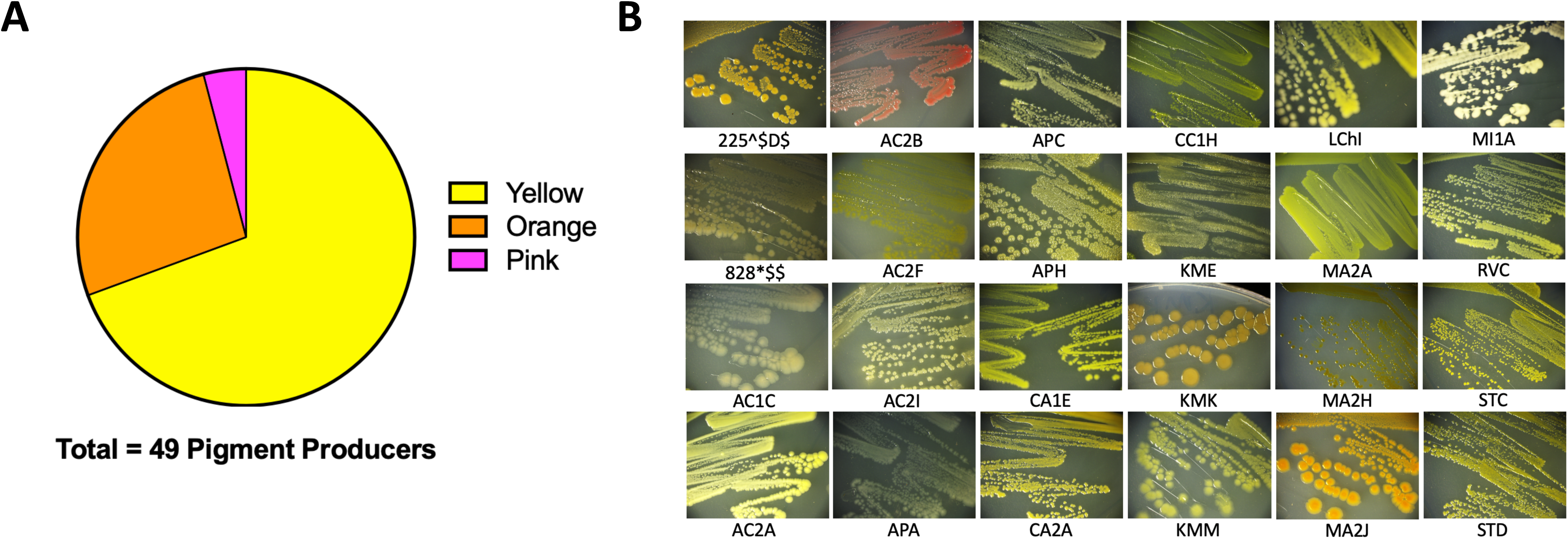
Extremophiles produce pigment. Pigment production was observed in colonies over seven days. (A) 49/142 colony isolates produce pigments. (B) Sample images of pigmented colonies after seven days of growth on R2 minimal medium agar at 25°C.

### Prominent biofilm formation is correlated to more extreme environments

Biofilms enhance colonization to surfaces, increase communication amongst cells, and enable symbioses (Vlamakis et al. 2013; Hall-Stoodley, Costerton, and Stoodley 2004) as well as protect against antimicrobials, DNA damage, mechanical forces, and other stresses (Hall-Stoodley, Costerton, and Stoodley 2004). To assess whether biofilm formation was used as a survival strategy in the extremophiles of our collection, we performed biofilm assays. Briefly, cells were inoculated into R2 medium and incubated statically for 5 days at 25°C in glass tubes. Using a standard Crystal Violet (CV) Assay, in which the CV dye stains cells that remain attached to surfaces, the degree of cell surface attachment was quantified by absorbance at OD_595_ of the resuspended CV (O’Toole 2011). Of all 142 isolates assayed under our conditions, 80% of isolates attached to surfaces (Figure 7). This was determined by any value greater or equal to 0.05AU, a value in which CV stained cells were visible on the glass. The species that had the greatest biofilm production across all isolates was *Bacillus simplex*.

**FIGURE 7.**
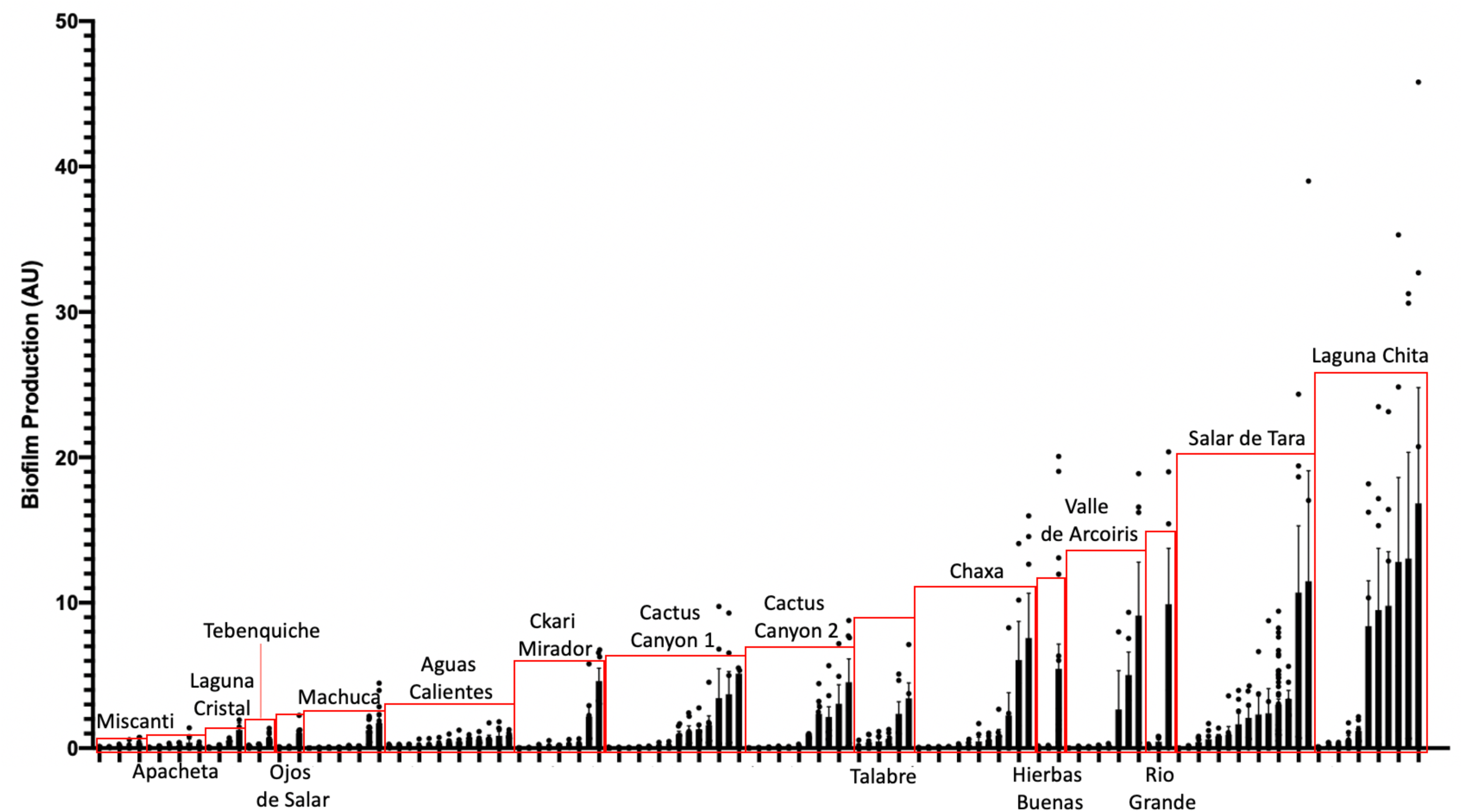
Extremophiles produce biofilm in more extreme environments. Colonies were assayed for biofilm development by inoculating colonies into R2 liquid medium in glass culture tubes. Cultures were grown statically at 25°C and quantified after 5 days of growth using the crystal violet (CV) staining assay (O’Toole 2011). Biofilms were stained with 0.1% CV. Solubilized CV was then quantified using a spectrophotometer at OD_595_. OD_595_ was standardized by cell count (OD_600_) to quantify the amount of biofilm produced per cell (AU). Bars represent the average of two or more independent experiments for each individual isolate, dots represent biological replicates (n≥6), and error bars represent standard deviation. Red boxes highlight isolates from the same location.

Also, of interest, were the isolates that produced a pellicle or clumped biofilm. Isolates that produced a visible pellicle, a biofilm formed at the air-liquid interface (Vlamakis et al. 2013), were KMK, KMM, KME, APH, AC2F, and STB (Figure S3). Additionally, KMK, RVC, AC2F, and APH produced a visible clumped biofilm while shaking. These are types of biofilms that might not be accurately quantified using the CV assay but are important examples of biofilm producers.

When comparing the amount of biofilm produced by isolates from the different locations, the ones from the most extreme environments were more likely to form biofilms. The bacteria inhabiting Laguna Chita had the most prominent biofilms; they attached to surfaces well. The location with the greatest number of biofilm producers was Salar de Tara (Figure 7). These locations are next to each other, are located at high elevation (>4200 meters above sea level), and are barren (Figure S1K and J, Table S1). The bacteria from these locations may use biofilms to facilitate nutrient sharing, protect against the high UV radiation, and enhance colonization. It is also of note that three isolates from Ckari Mirador produced pellicles/clumped biofilms. This location is exceptionally dry within the Atacama Desert and biofilm may be used by these isolates to enhance their survival in this extreme environment (Figure S1A).

Together, our data indicate that biofilm production is a strategy that bacteria use to survive in the extreme environments found in the Atacama Desert.

### Extremophiles produce antibiotics against Gram-positive and Gram-negative bacteria

From the 16S rRNA whole sample analysis, we found that 35.9% of the total bacteria in all the samples were *Actinobacteria* (Figure 8A). Within that phylum, 0.48% of all the total bacteria were within the genus *Streptomyces*. Talabre and Laguna Cristal (Figure S1G and L) had the highest relative abundance of *Streptomyces* at 4.6% and 1.5%, respectively (Table S2B). *Streptomyces* are known to contain many natural antibiotic compound producers (Barka et al. 2016). Since, we did not cultivate *Streptomyces*, we were curious to see if any of our non-*Streptomyces* isolates produced antibiotics. We chose a select group of isolates for careful investigation of antibiotic production. We selected the isolates from Tebenquiche, Machuca, and Hierbas Buenas (Figure S1O, C, and B), all outlier groups according to the 16S rRNA phylogenetic analysis (Figure 3C). Lastly, we had an interest in the *Enterobacter sp*. isolates being that *Enterobacter sp*. is on the list of ESKAPE pathogens (Santajit and Indrawattana 2016). This group of 21 selected isolates was assayed for antibiotic production (Table S4).

**FIGURE 8.**
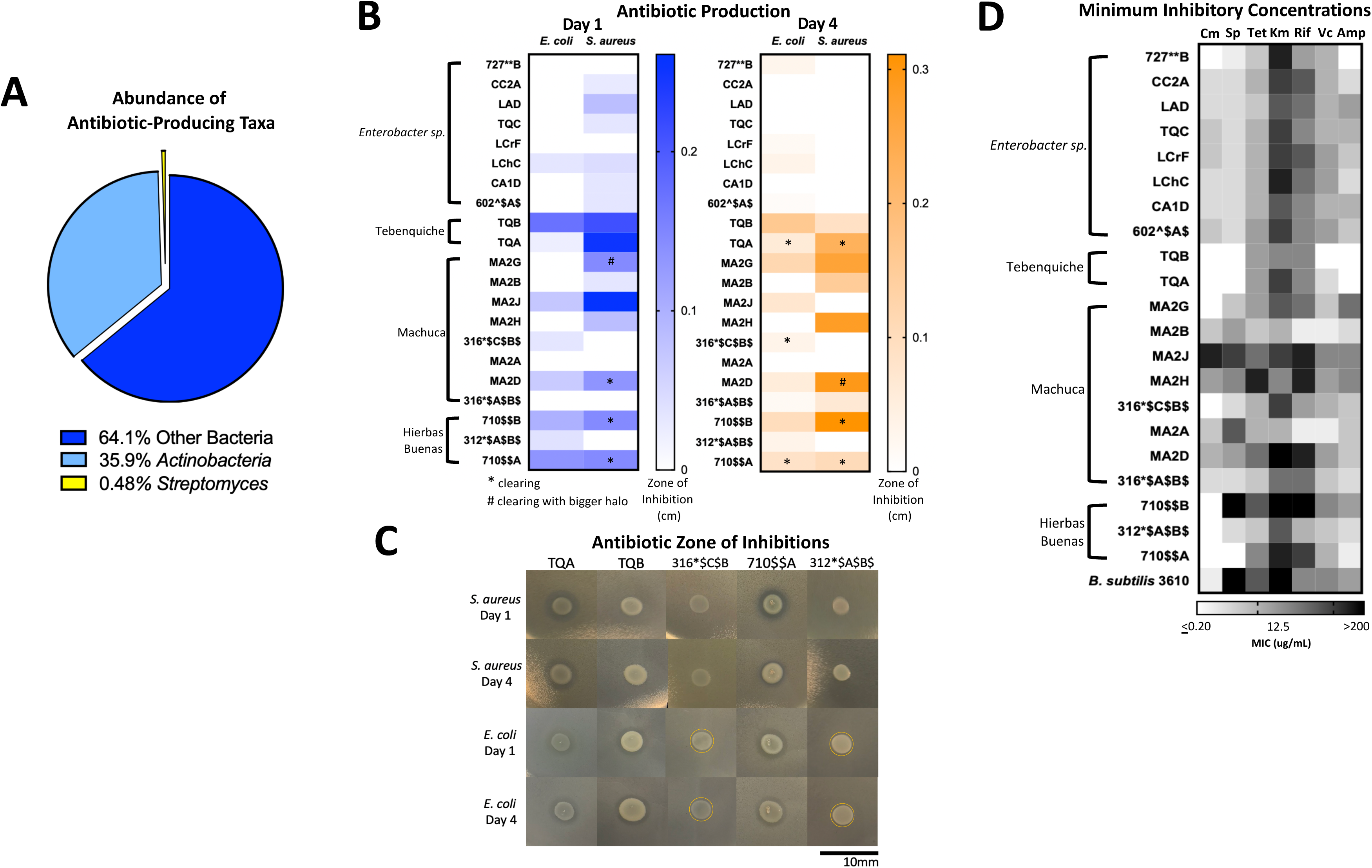
Extremophiles produce antimicrobials against *S. aureus* and *E. coli* and are naturally resistant to commercial antibiotics. (A) Abundance of potential antibiotic producing bacteria in total soil samples. Percent abundance of *Actinobacteria,* and its genus *Streptomyces* was calculated from the Illumina 16S sequencing data. (B and C) Antibiotic production assay demonstrated by antimicrobial challenge assay in which a select set of isolates inhibit the growth of either *S. aureus* or *E. coli* or both. Late stationary phase isolates that were not *Streptomyces* were spotted onto lawns of *S. aureus* and *E. coli.* (B) The zones of inhibition were measured after one and four days of incubation at 25°C. Stars indicate clearing and pound signs indicate clearing with a larger halo. Heatmap represents the average of three independent experiments each including three biological replicates (n=9). (C) Images of select isolates demonstrating antimicrobial production. Clearing can be seen in isolates TQA, TQB, and 710$$A. Yellow circle delineates haloing. (D) Minimum inhibition concentration (MIC) assay of select isolates demonstrated resistance to commercial antibiotics (Cm = chloramphenicol, Sp = spectinomycin, Tet = tetracycline, Km = kanamycin, Rif = rifampicin, Vc = vancomycin, Amp = ampicillin). In a 96-well plate, a select set of isolates were subjected to antibiotic challenge in 2-fold dilution in a range 200-0.2μg/mL. Cells were inoculated into R2 medium and grown statically at room temperature for 3 days. Growth was quantified by plate reader at OD600. The MIC was determined as the lowest concentration at which cell growth was completely inhibited. Heatmap represents the average MIC across three independent experiments in singlet (n=3).

To identify isolates that produced antibiotic, we performed a zone of inhibition assay against Gram-positive *Staphylococcus aureus* HG003 and Gram-negative *Escherichia coli* MG1655 as representatives of their respective groups. Isolates were grown shaking until late stationary phase and then spotted onto lawns of the target bacteria. Plates were incubated at 30°C and the zones of inhibitions were measured after one and four days of growth.

There were two types of inhibition phenomena; haloing and clearing (Figure 8C). Haloing was when there was a visible ring of inhibition, however the target bacteria still grew. Most of the isolates demonstrated this phenomenon. Possible explanations for this haloing could be a result of inhibition then gained resistance by the target bacteria, or due to different growth rates where the test isolate grew after the target bacterium resulting in a delayed killing. Regardless, the haloing demonstrated an inhibition of the target bacterium by the test isolate. Further analysis of the antibiotic resistant isolates would provide insights in the mechanism of resistance to the antibiotic. The other phenomenon, clearing, suggests the produced antibiotic is much more potent. The isolates that demonstrated clearing are indicated by stars or pound signs (Figure 8B) and the cleared zones can be seen in Figure 8C. Results show that many isolates produced antibiotics against *S. aureus* and *E. coli* (Figure 8B). 16 isolates inhibited the growth of *S. aureus,* while 13 exhibited a challenge to *E. coli.* A loss of inhibition after 4 days showed that the target bacteria grew resistant to the challenge.

Of the isolates, TQA (*Pseudomonas koreensis*, ID:98.32), TQB (*Pseudomonas azotoformans*, ID:99.26%) and 710$AA (*Pseudomonas brassicacearum*, ID: 99.7) produced the most potent inhibitory compounds against both *S. aureus* and *E. coli (*Figure 8B). This was seen by large clearing zones of inhibition (Figure 8C). Of more interest are isolates 316*$C$B$ (*Hafnia alvei*, ID: 98.1%), and 312*$A$B$ (*Hafnia alvei*, ID: 99.9%), both of which demonstrated killing against only the Gram-negative *E. coli* (Figure 8B). The identification of Gram-negative-specific antibiotic compounds are of particular interest for drug discovery groups (Lewis 2017).

Together, these data demonstrated that the chosen isolates produced antibiotics against *S. aureus* and *E. coli.* Antibiotic production is an expected strategy that provides competitive advantage in an extreme environment.

### Extremophiles are naturally resistant to commercial antibiotics

Coming from remote environmental samples, none of these isolates have been exposed to commercially available antibiotics. Many of these isolates were also identified as genera that have developed multi-drug resistances (Table S3, (Santajit and Indrawattana 2016)). Such examples include, *Enterobacter sp., Acinetobacter sp.,* and *Pseudomonas sp.* (Table S3). We were curious as to whether any of the isolates were inherently resistant to commercial antibiotics; thus, we performed a minimum inhibitory concentration (MIC) assay using antibiotics with different mechanisms of action. The MIC was measured for the same 21 isolates from Table S4 used in the antibiotic production assay. We determined the MIC for ampicillin (targets cell wall), vancomycin (targets cell wall), rifampicin (targets RNA polymerase), kanamycin (targets 30S ribosomal subunit), tetracycline (targets 30S ribosomal subunit), spectinomycin (targets 30S ribosomal subunit), and chloramphenicol (targets 50S ribosomal subunit) at a range of 200μg/mL to 0.2μg/mL (Lewis 2013). The MIC was defined as the lowest concentration of antibiotic that completely inhibited growth.

Results demonstrated that many isolates had high MICs to multiple antibiotics (Figure 8D). High MIC suggested resistance while low MIC suggested susceptibility to a particular antibiotic. Overall, the isolates had the highest MICs to ampicillin and the lowest MICs to kanamycin (Figure 8D, Table S5). The isolates with the highest MICs were TQB (*Pseudomonas azotoformans*: 99.26%) and MA2B (*Microbacterium sp*. ID: 99.2%) (Figure 8D, Table S3). For TQB, its lowest MIC was 6.25μg/mL against kanamycin and tetracycline (Table S5). TQB also had growth at 200μg/mL antibiotic concentration for four antibiotics (ampicillin, vancomycin, spectinomycin, and chloramphenicol) indicating it was not susceptible to these antibiotics at all (Table S5). For MA2B, its lowest MIC was 12.5μmL against vancomycin (Table S5). The most susceptible isolate was MA2J (*Exiguobacterium antarcticum* B7 ID:98.7%) (Figure 8D, Table S3). Its highest MIC was 6.25μg/mL against spectinomycin and chloramphenicol (Table S5). What gives us confidence for this assay is that isolates from the same genera had the same MIC patterning (Figure 8D). Examples include the *Enterobacter sp.* isolates (727**B*, CC2A, LAD, TQC, LCrF, LChC, CA1D, and 602^$A$) and the *Pseudomonas sp.* isolates (TQA, TQB, and 710$$A).

Together, these data demonstrate that isolates from remote extreme environments with little to no encounter with commercial antibiotics, have multidrug resistances.

## DISCUSSION

In this present study, we provide a comprehensive analysis of the microbial diversity using culture-independent and culture-dependent methods on environmental samples from 18 locations from within the Atacama Desert (Figures 1 to 4, Figures S1 and S2). Through the use of the more precise and wide-range method of Illumina 16S rRNA sequencing, we identified the dominant phyla to be *Actinobacteria* and *Proteobacteria* in Atacama soil (Figure 3B, Table S2). As expected, the locations with more plant life had a higher abundance of bacteria, and the two salt flat areas, Chaxa and Tebenquiche, were within the two outlier groups from the rest of the samples (Figure 3A and 3C, Figure S1E and S1O). A culture-dependent strategy and cultivated bacteria from the collected samples led to the isolation of 74/142 unique species, many of which were plant-associated (Figure 5, Table S3, Table 1).

Previous to this study, no one had reported the presence of bacteria in any of our locations except for Chaxa, Tebenquiche, and Miscanti (Figure S1E, O, H) (Dorador et al. 2018; Demergasso et al. 2004; Demergasso et al. 2008). These studies, however, were on water samples. Here, we add to the collection of data that others have provided to increase the understanding of the microbial diversity in new locations within the Atacama Desert.

We successfully isolated 74 unique species and a total of 142 isolates from all locations except for Baltinache (Figure SQ). We used direct cultivation in three media types as well as cultivation from a concentrated sample. These methods selected for aerobes that could grow at room temperature and that had the ability to survive transport over a month’s time. Our media was relatively rich, thus only those bacteria that could handle rich medium could grow. We also must keep in mind that our cultivation method was not designed to cultivate anaerobes, halophiles, acidophiles, or other condition-specific bacteria. Thus, the isolates that we cultivated must be extremotolerant bacteria; those that can tolerate extreme conditions but can also successfully grow under “normal” physiological conditions (Rampelotto 2013). Being that we cultivated under these conditions, it is highly probable that other bacteria can be cultivated from these samples using more selective conditions.

Many previous studies lacked research on bacteria physiology (Bull et al. 2016; Demergasso 2010; Demergasso et al. 2004; Demergasso et al. 2008; Mandakovic, Maldonado, et al. 2018; Mandakovic, Rojas, et al. 2018; Dorador et al. 2018; Dorador et al. 2010; Drees et al. 2006; Schulze-Makuch et al. 2018; Uritskiy et al. 2019). Being that a third of our isolates produced a pigment, there is strong likelihood that it is made for protection in its environment (Figure 6). One example of pigments used for survival is the production of pyocyanin and pyoveradine from *Pseudomonas sp*. Pyocyanin is a blue-green pigment produced by *Pseudomonas sp.* that acts an antimicrobial, but also has been shown to have siderophore activity in order to uptake environmental iron (Hassan and Fridovich 1980) (Jayaseelan, Ramaswamy, and Dharmaraj 2014). Pyoverdine, is a yellow-green pigment and siderophore secreted by *Pseudomonas sp.* during iron starvation as a way to sequester iron(III) (Meyer 2000). Indeed, the identified *Pseudomonas* genera isolates were seen to produce a secreted yellow pigment as exemplified by TQA, TQB, RGA, APA, and 710$$A (data not shown). Other groups have also seen that *Actinobacteria* and *Cyanobacteria* from the Atacama Desert produce melanins, scytonemin, carotenoids, chlorophylls, and other pigments (Azua-Bustos, Urrejola, and Vicuna 2012) (Bull and Asenjo 2013).

To date, no one has shown biofilm production data of microbial isolates from the Atacama Desert. 80% of our isolates produced a biofilm and there were many distinct types of biofilm produced (Figure 7 and Figure S3). The bacteria from these locations may use biofilm as a survival strategy to facilitate nutrient sharing, protect against the high UV index, and enhance colonization. The two locations of high biofilm production, Laguna Chita and Salar de Tara, were devoid of plant life (Figure S1K and J). The presence of biofilm-producing bacteria may also provide plant-promoting effects to the few low grasses that do live in the area.

Our study showed that some isolates produce antibiotics against Gram-positive *S. aureus* and Gram-negative *E. coli* (Figure 8). This was not surprising because the most dominant phylum was *Actinobacteria* (35.9%)(Rateb, Houssen, Arnold, et al. 2011; Rateb, Houssen, Harrison, et al. 2011; Undabarrena et al. 2016; Goodfellow et al. 2018; Okoro et al. 2009; Santhanam, Okoro, Rong, Huang, Bull, Andrews, et al. 2012; Santhanam, Okoro, Rong, Huang, Bull, Weon, et al. 2012) and it is likely other bacteria need to protect themselves against these. We found that the highest abundances of *Streptomyces* were in Talabre and Laguna Cristal (Table S2B, Figure S1G and L). Focusing on isolates from these locations will be important for finding novel antibiotic producers and compounds.

Lastly, we found that many isolates had high MICs against commercially produced antibiotics indicating inherent antibiotic resistance (Figure 8D). Being from remote environments, these isolates have theoretically not come in contact with commercial antibiotics until now. Because they are resistant to them, they must have intrinsic strategies to resist the antibiotic challenge. We believe this resistance has developed by three possible means: 1. they face antibiotic challenge in their environments, 2. mutagenesis as a result of being subjected to high levels of UV radiation, and 3. they are extremophiles and have strategies to withstand exogenous stress.

Many commercially available antibiotics were originally discovered to be produced by soil bacteria. Vancomycin, for example, was discovered from soil isolate *Streptomyces orientalis* (Levine 2006). It is, therefore, not surprising that our isolates had relatively high MICs to this antibiotic (Figure 8D). Environmental bacteria use antibiotics as a competitive advantage, and they must develop ways to resist those antibiotics and outcompete their challengers. Our isolates were most likely exposed to antibiotics in the soil and therefore, have developed resistance mechanisms that enable their growth when challenged with commercial antibiotics.

Mutagenesis is the result of a variety of genomic insults coming from diverse sources including reactive oxygen species, methylating agents, or ionizing radiation (Chatterjee and Walker 2017). UV radiation is one of the most damaging abiotic factors to the cell that directly damages the DNA by formation of thymidine dimers (Chatterjee and Walker 2017; Cordero et al. 2018). As a result, mutations occur, which could be favorable to the survival of bacterial species. Antibiotic resistance is one advantage that can be gained by mutagenesis (Li et al. 2019). Therefore, the more UV that the bacteria are exposed to, the higher the probability that they will accumulate mutations to confer antibiotic resistance.

The third explanation for higher resistance is that these extremophiles inherently have strategies to survive stress. These bacteria have improved their cell membranes, enzymes, DNA repair machineries, among other adaptations that allow them to resist chemical challenges in addition to the condition they already survive (Orellana et al. 2018). Efflux pumps, for example, have been found to increase antibiotic resistance and tolerance and are also found in many high metal tolerant extremophiles (Lewis 2017; Orellana et al. 2018). Such examples include *Arthrobacter sp.,* and *Flavobacterium sp.,* which were genera isolated and used in the MIC assay (Table S3, Table S4, Figure 8D) (Orellana et al. 2018). Studies have also shown that extremophiles from the Atacama Desert have high resistance to arsenic due to the arsenic-rich Atacama brooks and soils (Azua-Bustos, Urrejola, and Vicuna 2012; Escalante et al. 2009). Examples from our isolate list include *Pantoea, Serratia, Hafnia, Microbacterium, Exiguobacterium* and *Pseudomonas* (Tables S3 and S4) (Orellana et al. 2018). Indeed, our most resistant isolates were TQB (*Pseudomonas azotoformans*: 99.26%) and MA2B (*Microbacterium sp*. ID: 99.2%) (Figure 8D). Their resistance mechanisms for high metals and arsenic toxicity may lead to their naturally occurring antibiotic resistance.

### Conclusion

There are many reviews covering the advantages of extremophiles, especially in the light of bioremediation (Orellana et al. 2018; Azua-Bustos and Gonzalez-Silva 2014; Bull et al. 2016). It is important to go beyond the superficial microbial diversity analyses and isolate individual bacteria to learn what they are really doing. Cultivation is obviously a challenge but learning from extremophiles will allow us to understand the adaptation mechanisms that microbes use to survive in their environments. Their more sophisticated machineries will give insight into the metabolic and synthesis pathways that their non-extreme and pathogenic relatives utilize. This will lead to discoveries of new enzymes, pigments, and compounds that can be used in biotechnology, agriculture, and healthcare.

## EXPERIMENTAL PROCEDURES

### Sample Preparation and Storage

Environmental samples were collected during the months of May and June of 2018. At each location, soils were sampled from in between 1-5 cm and collected into sterile 2mL microcentrifuge tubes using sterile tools. Larger soil samples were also collected into sterile 50mL conical tubes. The date and exact coordinates were recorded at each sampling location (Table S1). It is important to note that samples from Chaxa, Miscanti, Aguas Calientes, and Salar de Tara (Figure S1E, H, I, J) were taken from outside of the national parks. Samples were maintained at room temperature while in the field. Once brought to the laboratory, 20% (v/v) glycerol in water was added to the samples and stored at −80° C.

### 16S Metagenomic Analysis

Approximately 1g of soil sample was added to UBiome microbiome sequencing kit tubes (UBiome, San Francisco, CA). Samples were processed and analyzed with the ZymoBIOMICS^®^ Service: Targeted Metagenomic Sequencing (Zymo Research, Irvine, CA).

#### Targeted Library Preparation

The ZymoBIOMICS^®^ DNA Miniprep Kit (Zymo Research, Irvine, CA) was used for DNA extraction. Bacterial 16S ribosomal RNA gene targeted sequencing was performed using the Quick-16S™ NGS Library Prep Kit (Zymo Research, Irvine, CA). The bacterial 16S primers amplified the V3-V4 region of the 16S rRNA gene using primers custom-designed by Zymo Research to provide the best coverage of the 16S gene while maintaining high sensitivity.

The sequencing library was prepared using a Zymo-developed library preparation process in which PCR reactions were performed in real-time PCR machines to control cycles and limited PCR chimera formation. The final PCR products were quantified with qPCR fluorescence readings and pooled together based on equal molarity. The final pooled library was cleaned up with the Select-a-Size DNA Clean & Concentrator™ (Zymo Research, Irvine, CA), then quantified with TapeStation^®^ and Qubit^®^.

#### Sequencing

The final library was sequenced on Illumina^®^ MiSeq™ with a v3 reagent kit (600 cycles). The sequencing was performed with >10% PhiX spike-in. Amplification was compared to a negative control standard.

#### Bioinformatics Analysis

Unique amplicon sequences were inferred from raw reads using the Dada2 pipeline (Callahan et al, 2016) (Callahan et al. 2016). Chimeric sequences were also removed with the Dada2 pipeline. Taxonomy was assigned using Uclust from Qiime v.1.9.1 with Greengenes 16S database as reference. Visualization and Analysis of Microbial Population Structure (VAMPS) program was used for all subsequent diversity, abundance, and cluster plot analyses (Huse et al. 2014). Alpha diversity, or richness, was quantified as the number of unique sequences found in the sample. Phyla and genera relative abundance were calculated as the percent number of sequences out of the total number of sequences in the sample. The cluster plots were calculated based on Morisita-Horn parameters at the genus level. For all analyses, the black soil sample was used as the representative soil sample from Valle de Arcoiris. This soil was chosen as it is the most prevalent soil type in the sampling location (Figure S1F).

### Bacteria Isolate Cultivation

Bacterial cultivation was performed using two approaches: direct cultivation and soil extract filter cultivation (Figure 2). For direct cultivation, ∼100 mg of soil was added directly to liquid cultivation medium. Four media were used: Luria Benton (LB) liquid and solid agar, 1:100 diluted LB liquid and solid agar, R2 liquid and solid agar (BD Difco, Franklin Lakes, NJ). Liquid cultures and plates were incubated at room temperature overnight. Liquid cultures that had visible growth, as seen by turbidity, were deposited onto the surface of solid agar plates of the same medium for colony formation. All cultures and plates were grown statically at room temperature.

The vacuum filter discs were collected to trap and concentrate any bacteria that were in the soil onto the filter paper. A sterile stick was used to scrape the filter disc and was resuspended into 100µL of sterile water. The resuspension was then plated onto R2 medium agar plates. Plates were incubated at room temperature for colony formation.

To isolate, colonies were struck onto R2 solid agar plates until single distinct colonies were identified. After obtaining all our isolates, potential sisters were removed by the following steps. First, colonies of the same morphology from the same location were removed. Second, colonies of the same morphology across the entire isolate collection were removed. At this point, isolates were stocked in 20% (v/v) glycerol in water and stored at −80°C.

For all subsequent cultivation and assays, R2 solid agar and liquid medium were used and incubated at 25°C. Isolates were observed to grow well in these conditions. Isolates were named according to a numbering system based on location, sample number, and medium type.

### Species Identification

To identify the species of each cultivated isolate we amplified and sequenced the 16S rRNA gene from the bacterial chromosome. A single colony was resuspended in 20µL of water and then used as the template in the polymerase chain reaction (PCR) reaction. The 16S rRNA gene was amplified using OneTaq Polymerase Master Mix (Fisher Scientific, Agawam, MA) and bacteria specific 16S rRNA gene primers 8F (5’-AGAGTTTGATCCTGGCTCAG-3’) and 1492R (5’-GGTTACCTTGTTACGACTT −3’) (Galkiewicz and Kellogg 2008). For those isolates that colony PCR was not successful, genomic DNA was extracted using chemical extraction and used as the template for PCR amplification (Sambrook and Russell 2001). The PCR products were sent for purification and Sanger Sequencing service at Eurofins Genomics (Louisville, Kentucky). Sequences were identified using NCBI Nucleotide Blast alignment.

We chose a 97% percent identity to determine species separation and confidence of the identified species. If the identity is lower than 97%, the isolate is considered a different and/or unidentified species. To identify unique isolates, we defined a sister isolate as any isolate different to another isolate within a given sample. If the percent identity was not available, then both isolates were kept. If both percent identities were above 97%, the isolate with the lower percent identity was removed. Cultivability ratio for each location was calculated by taking the number of its unique cultivated isolates (Figure 5A) and dividing it by its alpha diversity value (Figure 3A).

### Pigment Identification

Pigment production was observed by tracking morphology over the course of 7 days. Samples were categorized into yellow, orange, and pink pigment production. Results were reported as a percentage of all the isolates that produced a pigment.

### Biofilm Production Assay

Isolates were grown overnight in R2 liquid medium shaking at 25°C. Biofilm assay cultures were then inoculated 1:1000 into R2 liquid medium in small glass culture tubes and grown statically at room temperature. All strains were tested for biofilms after 5 days of static growth. At harvest, the OD600 was measured by spectrophotometer to quantify cell density. Quantification of biofilm production was done by adaptation of published protocols (O’Toole 2011). Briefly, cultures were decanted and rinsed with phosphate buffer saline (PBS). Any leftover liquid was pipetted out, and the tubes dried for 10 min in a fume hood. Each tube was filled with 0.1% (w/v) crystal violet stain and incubated for 15 min at room temperature. The stain was decanted, and the tubes were twice washed with PBS. Excess liquid was pipetted out and dried for 10 min under a flow hood. The remaining dye adhered to the biofilm was then dissolved in 30% (v/v) acetic acid in water. This solution solubilized the dye for 15 min. The crystal violet was measured at OD_595_. All experiments were performed in biological triplicate. R2 liquid medium was used as a blank control and the isolate 225*$C$, a known-biofilm producer, was used as a positive control in each experiment. For quantification of biofilm production, all OD_595_ values were subtracted by the blank control OD_595_ value. This new value was then divided by the isolate’s respective OD_600_ value to standardize all the cultures by cell count. Outliers were removed by Grubbs outlier test. Replicates were averaged, and the standard deviation of the mean was calculated to evaluate error.

### Antibiotic Production Assay

Test isolates were sub-cultured 1:100 from saturated cultures into R2 liquid medium in triplicate glass tubes. The cultures grew shaking at 25°C for 20 hrs. Lawns of *S. aureus* HG003 and *E. coli* MG1655 were prepared by sub-culturing cells 1:100 from saturated cultures into LB liquid medium. Bacterial cells were grown shaking at 30°C for one hour until reaching about OD_600_=0.1, which were then diluted to an OD_600_=0.02 in R2 liquid medium. The diluted cells were used to flood R2 solid agar plates and further dried in a flow hood for 30min. 3µL of the environmental isolates’ saturated cultures were spotted onto the *E. coli* and *S. aureus* lawns. For comparison purposes, ampicillin (25mg/mL) was spotted as a positive control since the strains are ampicillin susceptible, and R2 liquid medium was spotted as a negative control. Plates were incubated at 25°C. After one and four days of incubation, the zones of inhibition were measured from the edge of the grown colony to the edge of the inhibition zone. Three independent experiments each with biological triplicates were performed. The average zone of inhibition was calculated for all three experiments (n=9).

### Minimum Inhibitory Concentration (MIC) Dilution Assay

Selected isolates were challenged against commercial antibiotics ampicillin, rifampicin, vancomycin, kanamycin, tetracycline, spectinomycin, and chloramphenicol (all purchased from Fisher Scientific, Agawam, MA or Sigma Millipore, St. Louis, MO) in 96-well plates. Media was supplemented with antibiotic in a 2-fold dilution ranging from 200 μg/mL to 0.2 μg/mL final concentration. Rifampicin was diluted 2-fold in a range 164 μg/mL to 0.2 μg/mL final concentration. Cells were inoculated at a 1:100 dilution from a saturated culture. Wells with no antibiotic, no cells, and R2 supplemented with antibiotic vehicle were included as positive, negative, and vehicle effect controls. *Bacillus subtilis* NCBI 3610 was used as a control strain with known MICs to these antibiotics. Assay plates were covered with a sealing film and incubated at room temperature for three days. After three days, isolate growth was quantified using a plate reader at OD600. The MIC was determined as the concentration that fully inhibited growth. The experiment was performed three independent times each in biological singlet. The MIC was calculated to be the average from the three experiments.

## ACKNOWLEDGEMENTS

The authors would like to thank Jonna Iacono, Director of Office of Undergraduate Research and Fellowships, at Northeastern University for co-leading the trips to Chile. The authors are thankful to the Northeastern University undergraduate students that participated in the Dialogue of Civilizations courses to Chile (2017&2018). The students performed sample collection, bacterial growth and colony purification, DNA isolation, and PCR amplifications while in Chile as part of the course. We would also like to thank Joey Lehman Morris for being an active participant in the science aspect of the course while teaching students about the art of landscape photography. OneSeed Expeditions for organizing the trip in Chile. Special thanks to our guide Sofia Mardones and to our most patient and knowledgeable driver Guillermo Maluenda. We would also like to thank students Paul Mueller and Hannah Meiseles for bioinformatics assistance, and the students at University of Antofagasta for lab support while in Chile. Thank you to Dr. Slava Epstein from Northeastern University Department of Biology and Dr. D. Mark Welch from Marine Biological Laboratory Woods Hole, MA for insightful discussion and revisions of the paper. Finally, thanks to the Godoy-Carter and Chai Lab members for continuous discussion and support. This work was funded by Northeastern University (NU) Global Experience Office, and the NU Scholars Programs. V.G.G is funded by a stipend from NuSci, an Inclusive Excellence grant from HHMI, Y. C. is supported by a National Science Foundation grant (MCB1651732), and A.R. is supported by the NU Provost Dissertation Completion Fellowship.

## SUPPORTING INFORMATION

**Table S1.** Location characteristics

**Table S2.** Percent relative abundance of phyla and genera in all samples

**Table S3.** List of cultivated isolates

**Table S4.** List of select isolate for MIC and antibiotic production assays

**Table S5.** Select isolate MIC values

**FIGURE S1.**
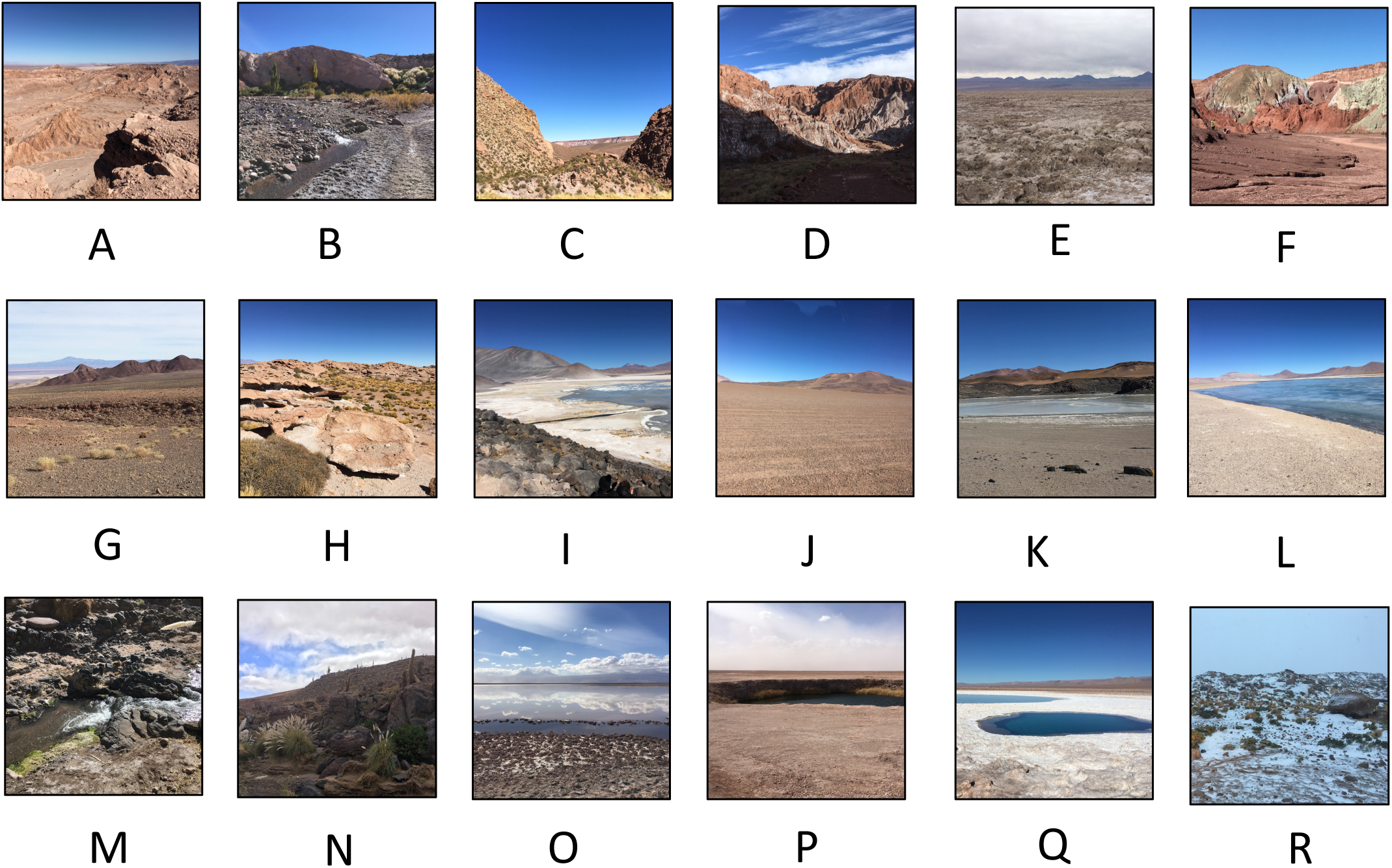
Images of sampling locations. Images taken at the time of sampling in each location. Letter corresponds to the letter identifier in Figure 1.

**FIGURE S2.**
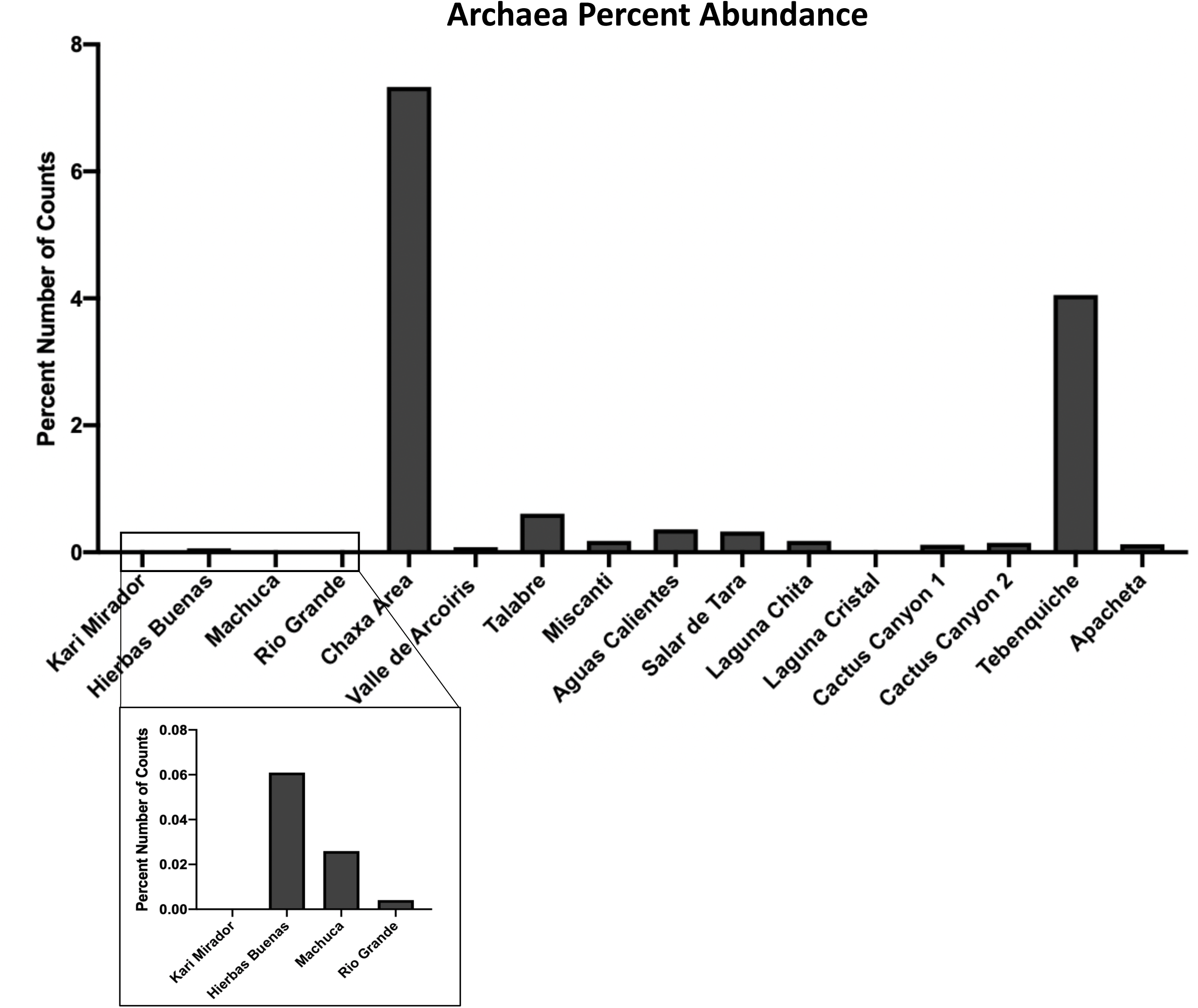
Archaea make up 1.14% of the total community diversity. The percent number of Archaea sequences of total samples was calculated from the Illumina 16S rRNA sequencing data set for each location. The total makeup of all detected sequences was 1.14% Archaea. Chaxa and Tebenquiche had the highest abundance of Archaea. Ckari Mirador and Laguna Cristal contained no detection of Archaea (box is a zoomed in look at the values).

**FIGURE S3.**
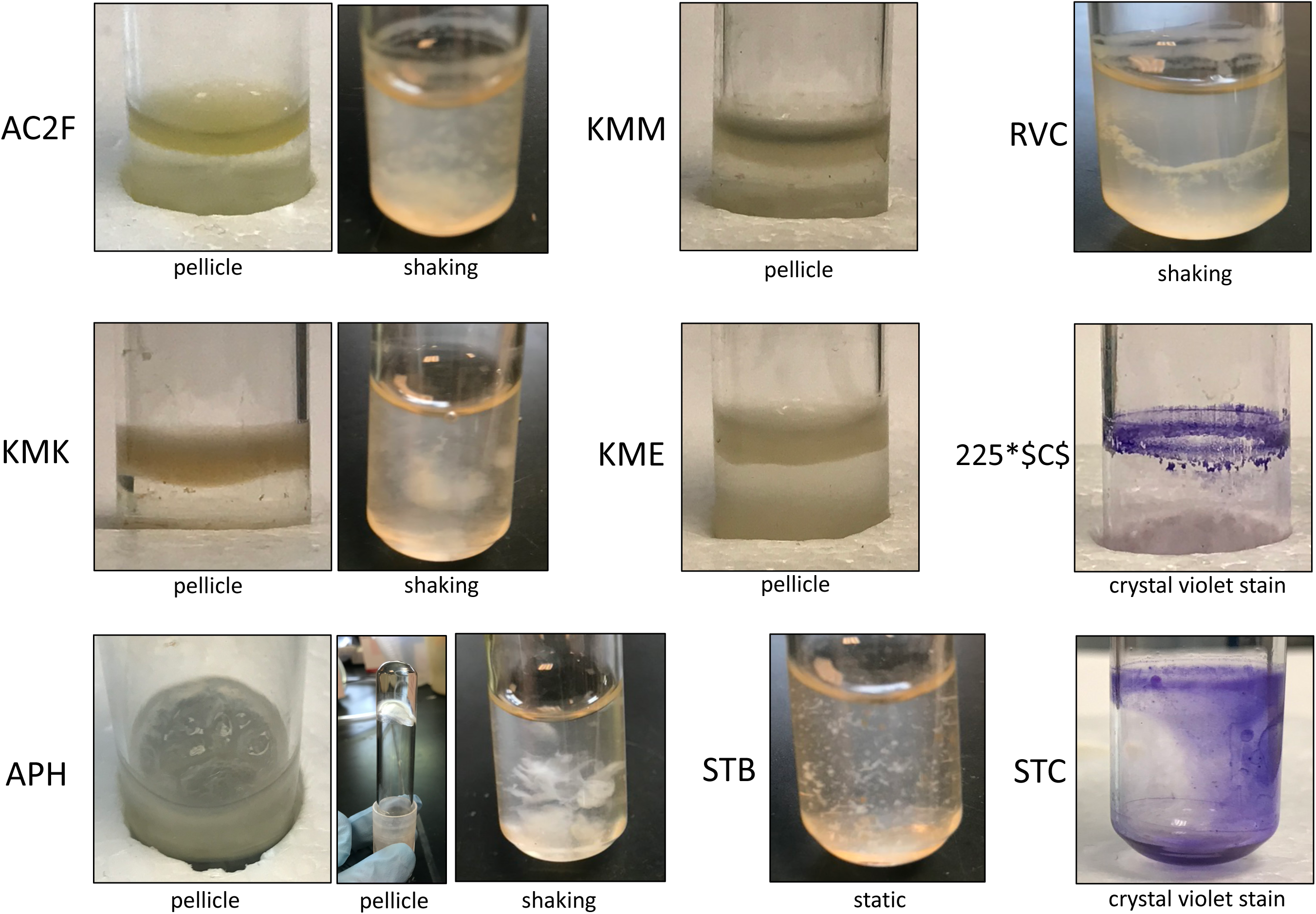
Images of unique biofilm producers. Side view images taken of unique biofilm producers. Pellicle biofilms were produced after 5 days of static incubation at 25°C in R2 liquid medium. Shaking biofilms were observed as excess matrix and cell clumping. Cells were grown overnight shaking at 25°C in R2 liquid medium. Crystal violet-stained biofilms were imaged after 5 days of static incubation at 25°C in R2 liquid medium.

## REFERENCES

Aguayo, P., P. Gonzalez, V. Campos, T. L. Maugeri, M. Papale, C. Gugliandolo, and M. A. Martinez. 2017. ‘Comparison of Prokaryotic Diversity in Cold, Oligotrophic Remote Lakes of Chilean Patagonia’, Curr Microbiol, 74: 598–613.

Anda, M., Y. Ohtsubo, T. Okubo, M. Sugawara, Y. Nagata, M. Tsuda, K. Minamisawa, and H. Mitsui. 2015. ‘Bacterial clade with the ribosomal RNA operon on a small plasmid rather than the chromosome’, Proc Natl Acad Sci U S A, 112: 14343–7.

Azua-Bustos, A., L. Caro-Lara, and R. Vicuna. 2015. ‘Discovery and microbial content of the driest site of the hyperarid Atacama Desert, Chile’, Environ Microbiol Rep, 7: 388–94.

Azua-Bustos, A., and C. Gonzalez-Silva. 2014. ‘Biotechnological applications derived from microorganisms of the Atacama Desert’, Biomed Res Int, 2014: 909312.

Azua-Bustos, A., C. Urrejola, and R. Vicuna. 2012. ‘Life at the dry edge: microorganisms of the Atacama Desert’, FEBS Lett, 586: 2939–45.

Barka, E. A., P. Vatsa, L. Sanchez, N. Gaveau-Vaillant, C. Jacquard, J. P. Meier-Kolthoff, H. P. Klenk, C. Clement, Y. Ouhdouch, and G. P. van Wezel. 2016. ‘Taxonomy, Physiology, and Natural Products of Actinobacteria’, Microbiol Mol Biol Rev, 80: 1–43.

Bates, S. T., D. Berg-Lyons, J. G. Caporaso, W. A. Walters, R. Knight, and N. Fierer. 2011. ‘Examining the global distribution of dominant archaeal populations in soil’, ISME J, 5: 908–17.

Beaz-Hidalgo, R., F. Latif-Eugenin, M. J. Hossain, K. Berg, R. M. Niemi, J. Rapala, C. Lyra, M. R. Liles, and M. J. Figueras. 2015. ‘Aeromonas aquatica sp. nov., Aeromonas finlandiensis sp. nov. and Aeromonas lacus sp. nov. isolated from Finnish waters associated with cyanobacterial blooms’, Syst Appl Microbiol, 38: 161–8.

Berdy, B., A. L. Spoering, L. L. Ling, and S. S. Epstein. 2017. ‘In situ cultivation of previously uncultivable microorganisms using the ichip’, Nat Protoc, 12: 2232–42.

Bull, A. T., B. A. Andrews, C. Dorador, and M. Goodfellow. 2018. ‘Introducing the Atacama Desert’, Antonie Van Leeuwenhoek, 111: 1269–72.

Bull, A. T., and J. A. Asenjo. 2013. ‘Microbiology of hyper-arid environments: recent insights from the Atacama Desert, Chile’, Antonie Van Leeuwenhoek, 103: 1173–9.

Bull, A. T., J. A. Asenjo, M. Goodfellow, and B. Gomez-Silva. 2016. ‘The Atacama Desert: Technical Resources and the Growing Importance of Novel Microbial Diversity’, Annu Rev Microbiol, 70: 215–34.

Callahan, B. J., P. J. McMurdie, M. J. Rosen, A. W. Han, A. J. Johnson, and S. P. Holmes. 2016. ‘DADA2: High-resolution sample inference from Illumina amplicon data’, Nat Methods, 13: 581–3.

Caneschi, W. L., A. B. Sanchez, E. B. Felestrino, C. G. C. Lemes, I. F. Cordeiro, N. P. Fonseca, M. M. Villa, I. T. Vieira, L. A. G. Moraes, R. A. B. Assis, F. F. do Carmo, L. H. Y. Kamino, R. S. Silva, J. A. Ferro, M. I. T. Ferro, R. M. Ferreira, V. L. Santos, U. C. M. Silva, N. F. Almeida, A. M. Varani, C. C. M. Garcia, J. C. Setubal, and L. M. Moreira. 2019. ‘Serratia liquefaciens FG3 isolated from a metallophyte plant sheds light on the evolution and mechanisms of adaptive traits in extreme environments’, Sci Rep, 9: 18006.

Chatterjee, N., and G. C. Walker. 2017. ‘Mechanisms of DNA damage, repair, and mutagenesis’, Environ Mol Mutagen, 58: 235–63.

Chen, Q., W. A. Meyer, Q. Zhang, and J. F. White. 2020. ‘16S rRNA metagenomic analysis of the bacterial community associated with turf grass seeds from low moisture and high moisture climates’, PeerJ, 8: e8417.

Cordero, R. R., A. Damiani, J. Jorquera, E. Sepulveda, M. Caballero, S. Fernandez, S. Feron, P. J. Llanillo, J. Carrasco, D. Laroze, and F. Labbe. 2018. ‘Ultraviolet radiation in the Atacama Desert’, Antonie Van Leeuwenhoek, 111: 1301–13.

D’Argenio, V., and F. Salvatore. 2015. ‘The role of the gut microbiome in the healthy adult status’, Clin Chim Acta, 451: 97–102.

de Carvalho, Cccr. 2017. ‘Biofilms: Microbial Strategies for Surviving UV Exposure’, Adv Exp Med Biol, 996: 233–39.

Deb, S., and S. K. Das. 2020. ‘Draft Genome Sequence of Microbacterium oryzae Strain MB-10, Isolated from a Rice Field in India’, Microbiol Resour Announc, 9.

Demergasso, C., E. O. Casamayor, G. Chong, P. Galleguillos, L. Escudero, and C. Pedros-Alio. 2004. ‘Distribution of prokaryotic genetic diversity in athalassohaline lakes of the Atacama Desert, Northern Chile’, FEMS Microbiol Ecol, 48: 57–69.

Demergasso, C., Dorador, C., Meneses, D., Blamey, J., Cabrol, N., Escudero, L., Chong, G. 2010. ‘Prokaryotic diversity pattern in high-altitude ecosystems of the Chilean Altiplano’, Journal of Geophysical Research, 115.

Demergasso, C., L. Escudero, E. O. Casamayor, G. Chong, V. Balague, and C. Pedros-Alio. 2008. ’Novelty and spatio-temporal heterogeneity in the bacterial diversity of hypersaline Lake Tebenquiche (Salar de Atacama)’, Extremophiles, 12: 491–504.

Dorador, C., P. Fink, M. Hengst, G. Icaza, A. S. Villalobos, D. Vejar, D. Meneses, V. Zadjelovic, L. Burmann, J. Moelzner, and C. Harrod. 2018. ‘Microbial community composition and trophic role along a marked salinity gradient in Laguna Puilar, Salar de Atacama, Chile’, Antonie Van Leeuwenhoek, 111: 1361–74.

Dorador, C., I. Vila, F. Remonsellez, J. F. Imhoff, and K. P. Witzel. 2010. ‘Unique clusters of Archaea in Salar de Huasco, an athalassohaline evaporitic basin of the Chilean Altiplano’, FEMS Microbiol Ecol, 73: 291–302.

Drees, K. P., J. W. Neilson, J. L. Betancourt, J. Quade, D. A. Henderson, B. M. Pryor, and R. M. Maier. 2006. ‘Bacterial community structure in the hyperarid core of the Atacama Desert, Chile’, Appl Environ Microbiol, 72: 7902–8.

Escalante, G., V. L. Campos, C. Valenzuela, J. Yanez, C. Zaror, and M. A. Mondaca. 2009. ‘Arsenic resistant bacteria isolated from arsenic contaminated river in the Atacama Desert (Chile)’, Bull Environ Contam Toxicol, 83: 657–61.

Escudero L, Chong G, Demergasso C, Farias ME, Cabrol NA, Grin E, Minkley Jr. E, Yu Y. 2007. ‘Investigating microbial diversity and UV radiation impact at the high-altitude Lake Aguas Calientes, Chile’, Proc, SPIE 6694 Instruments, Methods, and Missions for Astrobiology X, 66940Z.

Fang, Y., L. Wu, G. Chen, and G. Feng. 2016. ‘Complete genome sequence of Pseudomonas azotoformans S4, a potential biocontrol bacterium’, J Biotechnol, 227: 25–26.

Ferri, M., E. Ranucci, P. Romagnoli, and V. Giaccone. 2017. ‘Antimicrobial resistance: A global emerging threat to public health systems’, Crit Rev Food Sci Nutr, 57: 2857–76.

Fu, Y., Y. Wu, Y. Yuan, and M. Gao. 2017. ‘Complete Genome Sequence of Bacillus thuringiensis Serovar rongseni Reference Strain SCG04-02, a Strain Toxic to Plutella xylostella’, Genome Announc, 5.

Galkiewicz, J. P., and C. A. Kellogg. 2008. ‘Cross-kingdom amplification using bacteria-specific primers: complications for studies of coral microbial ecology’, Appl Environ Microbiol, 74: 7828–31.

George, P. B. L., D. Lallias, S. Creer, F. M. Seaton, J. G. Kenny, R. M. Eccles, R. I. Griffiths, I. Lebron, B. A. Emmett, D. A. Robinson, and D. L. Jones. 2019. ‘Divergent national-scale trends of microbial and animal biodiversity revealed across diverse temperate soil ecosystems’, Nat Commun, 10: 1107.

Gerardin, Y., M. Springer, and R. Kishony. 2016. ‘A competitive trade-off limits the selective advantage of increased antibiotic production’, Nat Microbiol, 1: 16175.

Goodfellow, M., I. Nouioui, R. Sanderson, F. Xie, and A. T. Bull. 2018. ‘Rare taxa and dark microbial matter: novel bioactive actinobacteria abound in Atacama Desert soils’, Antonie Van Leeuwenhoek, 111: 1315–32.

Graupner, K., G. Lackner, and C. Hertweck. 2015. ‘Genome Sequence of Mushroom Soft-Rot Pathogen Janthinobacterium agaricidamnosum’, Genome Announc, 3.

Guo, Y., Z. Jiao, L. Li, D. Wu, D. E. Crowley, Y. Wang, and W. Wu. 2012. ‘Draft genome sequence of Rahnella aquatilis strain HX2, a plant growth-promoting rhizobacterium isolated from vineyard soil in Beijing, China’, J Bacteriol, 194: 6646–7.

Gutiérrez-Luna, F.M., López-Bucio, J., Altamirano-Hernández, J., Valencia-Cantero, E., Reyes de la Cruz, H., and Macías-Rodríguez, L. 2010. ‘Plant growth-promoting rhizobacteria modulate root-system architecture in Arabidopsis thaliana through volatile organic compound emission’, Symbiosis, 51: 75–83.

Hall, C. W., and T. F. Mah. 2017. ‘Molecular mechanisms of biofilm-based antibiotic resistance and tolerance in pathogenic bacteria’, FEMS Microbiol Rev, 41: 276–301.

Hall-Stoodley, L., J. W. Costerton, and P. Stoodley. 2004. ‘Bacterial biofilms: from the natural environment to infectious diseases’, Nat Rev Microbiol, 2: 95–108.

Hassan, H. M., and I. Fridovich. 1980. ‘Mechanism of the antibiotic action pyocyanine’, J Bacteriol, 141: 156–63.

He, A. L., S. Q. Niu, Q. Zhao, Y. S. Li, J. Y. Gou, H. J. Gao, S. Z. Suo, and J. L. Zhang. 2018. ‘Induced Salt Tolerance of Perennial Ryegrass by a Novel Bacterium Strain from the Rhizosphere of a Desert Shrub Haloxylon ammodendron’, Int J Mol Sci, 19.

Huang, F. L., Y. Zhang, L. P. Zhang, S. Wang, Y. Feng, and N. H. Rong. 2019. ‘Complete genome sequence of Bacillus megaterium JX285 isolated from Camellia oleifera rhizosphere’, Comput Biol Chem, 79: 1–5.

Huse, S. M., D. B. Mark Welch, A. Voorhis, A. Shipunova, H. G. Morrison, A. M. Eren, and M. L. Sogin. 2014. ‘VAMPS: a website for visualization and analysis of microbial population structures’, BMC Bioinformatics, 15: 41.

Jayaseelan, S., D. Ramaswamy, and S. Dharmaraj. 2014. ‘Pyocyanin: production, applications, challenges and new insights’, World J Microbiol Biotechnol, 30: 1159–68.

Jeong, H. W., M. S. Bang, Y. J. Lee, S. J. Lee, S. C. Lee, J. I. Shin, and C. H. Oh. 2018. ‘Complete Genome Sequence of Bacillus subtilis Strain DKU_NT_03, Isolated from a Traditional Korean Food Using Soybean (Chung-gook-jang) for High-Quality Nattokinase Activity’, Genome Announc, 6.

Jung, B. K., A. R. Khan, S. J. Hong, G. S. Park, Y. J. Park, C. E. Park, H. J. Jeon, S. E. Lee, and J. H. Shin. 2017. ‘Genomic and phenotypic analyses of Serratia fonticola strain GS2: a rhizobacterium isolated from sesame rhizosphere that promotes plant growth and produces N-acyl homoserine lactone’, J Biotechnol, 241: 158–62.

Kerrigan, Z., J. B. Kirkpatrick, and S. D’Hondt. 2019. ‘Influence of 16S rRNA Hypervariable Region on Estimates of Bacterial Diversity and Community Composition in Seawater and Marine Sediment’, Front Microbiol, 10: 1640.

Labeda, D. P. 2001. ‘Crossiella gen. nov., a new genus related to Streptoalloteichus’, Int J Syst Evol Microbiol, 51: 1575–79.

Levine, D. P. 2006. ’Vancomycin: a history’, *Clin Infect Dis*, 42 Suppl 1: S5-12. Lewis, K. 2013. ‘Platforms for antibiotic discovery’, Nat Rev Drug Discov, 12: 371–87.

Levine, D. P.. 2017. ‘New approaches to antimicrobial discovery’, Biochem Pharmacol, 134: 87–98.

Li, X., A. Z. Gu, Y. Zhang, B. Xie, D. Li, and J. Chen. 2019. ‘Sub-lethal concentrations of heavy metals induce antibiotic resistance via mutagenesis’, J Hazard Mater, 369: 9–16.

Lin, H., S. Hu, R. Liu, P. Chen, C. Ge, B. Zhu, and L. Guo. 2016. ‘Genome Sequence of Pseudomonas koreensis CRS05-R5, an Antagonistic Bacterium Isolated from Rice Paddy Field’, Front Microbiol, 7: 1756.

Ma, J., C. Wang, H. Wang, K. Liu, T. Zhang, L. Yao, Z. Zhao, B. Du, and Y. Ding. 2018. ‘Analysis of the Complete Genome Sequence of Bacillus atrophaeus GQJK17 Reveals Its Biocontrol Characteristics as a Plant Growth-Promoting Rhizobacterium’, Biomed Res Int, 2018: 9473542.

Mandakovic, D., J. Maldonado, R. Pulgar, P. Cabrera, A. Gaete, V. Urtuvia, M. Seeger, V. Cambiazo, and M. Gonzalez. 2018. ‘Microbiome analysis and bacterial isolation from Lejia Lake soil in Atacama Desert’, Extremophiles, 22: 665–73.

Mandakovic, D., C. Rojas, J. Maldonado, M. Latorre, D. Travisany, E. Delage, A. Bihouee, G. Jean, F. P. Diaz, B. Fernandez-Gomez, P. Cabrera, A. Gaete, C. Latorre, R. A. Gutierrez, A. Maass, V. Cambiazo, S. A. Navarrete, D. Eveillard, and M. Gonzalez. 2018. ‘Structure and co-occurrence patterns in microbial communities under acute environmental stress reveal ecological factors fostering resilience’, Sci Rep, 8: 5875.

Meyer, J. M. 2000. ‘Pyoverdines: pigments, siderophores and potential taxonomic markers of fluorescent Pseudomonas species’, Arch Microbiol, 174: 135–42.

Narsing Rao, M. P., M. Xiao, and W. J. Li. 2017. ‘Fungal and Bacterial Pigments: Secondary Metabolites with Wide Applications’, Front Microbiol, 8: 1113.

O’Toole, G. A. 2011. ‘Microtiter dish biofilm formation assay’, J Vis Exp.

Okoro, C. K., R. Brown, A. L. Jones, B. A. Andrews, J. A. Asenjo, M. Goodfellow, and A. T. Bull. 2009. ‘Diversity of culturable actinomycetes in hyper-arid soils of the Atacama Desert, Chile’, Antonie Van Leeuwenhoek, 95: 121–33.

Ordenes-Aenishanslins, N., G. Anziani-Ostuni, M. Vargas-Reyes, J. Alarcon, A. Tello, and J. M. Perez-Donoso. 2016. ‘Pigments from UV-resistant Antarctic bacteria as photosensitizers in Dye Sensitized Solar Cells’, J Photochem Photobiol B, 162: 707–14.

Orellana, R., C. Macaya, G. Bravo, F. Dorochesi, A. Cumsille, R. Valencia, C. Rojas, and M. Seeger. 2018. ‘Living at the Frontiers of Life: Extremophiles in Chile and Their Potential for Bioremediation’, Front Microbiol, 9: 2309.

Oren, A. 2013. ‘Salinibacter: an extremely halophilic bacterium with archaeal properties’, FEMS Microbiol Lett, 342: 1–9.

Parro, V., G. de Diego-Castilla, M. Moreno-Paz, Y. Blanco, P. Cruz-Gil, J. A. Rodriguez-Manfredi, D. Fernandez-Remolar, F. Gomez, M. J. Gomez, L. A. Rivas, C. Demergasso, A. Echeverria, V. N. Urtuvia, M. Ruiz-Bermejo, M. Garcia-Villadangos, M. Postigo, M. Sanchez-Roman, G. Chong-Diaz, and J. Gomez-Elvira. 2011. ‘A microbial oasis in the hypersaline Atacama subsurface discovered by a life detector chip: implications for the search for life on Mars’, Astrobiology, 11: 969–96.

Ramette, A., M. Frapolli, M. Fischer-Le Saux, C. Gruffaz, J. M. Meyer, G. Defago, L. Sutra, and Y. Moenne-Loccoz. 2011. ‘Pseudomonas protegens sp. nov., widespread plant-protecting bacteria producing the biocontrol compounds 2,4-diacetylphloroglucinol and pyoluteorin’, Syst Appl Microbiol, 34: 180–8.

Rampelotto, P. H. 2013. ‘Extremophiles and extreme environments’, Life (Basel), 3: 482–5.

Rateb, M. E., W. E. Houssen, M. Arnold, M. H. Abdelrahman, H. Deng, W. T. Harrison, C. K. Okoro, J. A. Asenjo, B. A. Andrews, G. Ferguson, A. T. Bull, M. Goodfellow, R. Ebel, and M. Jaspars. 2011. ‘Chaxamycins A-D, bioactive ansamycins from a hyper-arid desert Streptomyces sp’, J Nat Prod, 74: 1491–9.

Rateb, M. E., W. E. Houssen, W. T. Harrison, H. Deng, C. K. Okoro, J. A. Asenjo, B. A. Andrews, A. T. Bull, M. Goodfellow, R. Ebel, and M. Jaspars. 2011. ‘Diverse metabolic profiles of a Streptomyces strain isolated from a hyper-arid environment’, J Nat Prod, 74: 1965–71.

Rundel P, Villagra P, Dillon MC, Roig S, and Debandi G. 2007. ‘Deserts and semi-desert environments’, Oxford Regional Environment Series: 153–83.

S, Jeyanthi V and Kanimozhi. 2018. ‘Plant Growth Promoting Rhizobacteria (PGPR) – Prospective and Mechanisms: A Review’, Journal of Pure and Applied Microbiology, 12: 733–49.

Sambrook, Joseph, and David W. Russell. 2001. Molecular cloning : a laboratory manual (Cold Spring Harbor Laboratory Press: Cold Spring Harbor, N.Y.).

Santajit, S., and N. Indrawattana. 2016. ‘Mechanisms of Antimicrobial Resistance in ESKAPE Pathogens’, Biomed Res Int, 2016: 2475067.

Santhanam, R., C. K. Okoro, X. Rong, Y. Huang, A. T. Bull, B. A. Andrews, J. A. Asenjo, H. Y. Weon, and M. Goodfellow. 2012. ‘Streptomyces deserti sp. nov., isolated from hyper-arid Atacama Desert soil’, Antonie Van Leeuwenhoek, 101: 575–81.

Santhanam, R., C. K. Okoro, X. Rong, Y. Huang, A. T. Bull, H. Y. Weon, B. A. Andrews, J. A. Asenjo, and M. Goodfellow. 2012. ‘Streptomyces atacamensis sp. nov., isolated from an extreme hyper-arid soil of the Atacama Desert, Chile’, Int J Syst Evol Microbiol, 62: 2680–4.

Schulze-Makuch, D., D. Wagner, S. P. Kounaves, K. Mangelsdorf, K. G. Devine, J. P. de Vera, P. Schmitt-Kopplin, H. P. Grossart, V. Parro, M. Kaupenjohann, A. Galy, B. Schneider, A. Airo, J. Frosler, A. F. Davila, F. L. Arens, L. Caceres, F. S. Cornejo, D. Carrizo, L. Dartnell, J. DiRuggiero, M. Flury, L. Ganzert, M. O. Gessner, P. Grathwohl, L. Guan, J. Heinz, M. Hess, F. Keppler, D. Maus, C. P. McKay, R. U. Meckenstock, W. Montgomery, E. A. Oberlin, A. J. Probst, J. S. Saenz, T. Sattler, J. Schirmack, M. A. Sephton, M. Schloter, J. Uhl, B. Valenzuela, G. Vestergaard, L. Wormer, and P. Zamorano. 2018. ‘Transitory microbial habitat in the hyperarid Atacama Desert’, Proc Natl Acad Sci U S A, 115: 2670–75.

Sears, C. L. 2005. ‘A dynamic partnership: celebrating our gut flora’, Anaerobe, 11: 247–51.

See-Too, W. S., Y. L. Lim, R. Ee, P. Convey, D. A. Pearce, W. F. Yin, and K. G. Chan. ‘Complete genome of Pseudomonas sp. strain L10.10, a psychrotolerant biofertilizer that could promote plant growth.’, Journal of Biotechnology, 222: 84–85.

See-Too, W.S., Convey, P., Pearce, D.A., Lim, Y.L., Ee R., Yin W.F., and Chan, K.G. 2016. ‘Complete genome of Planococcus rifietoensis M8(T), a halotolerant and potentially plant growth promoting bacterium’, Journal of Biotechnology, 221: 114–5.

Seki, T., A. Matsumoto, S. Omura, and Y. Takahashi. 2015. ‘Distribution and isolation of strains belonging to the order Solirubrobacterales’, J Antibiot (Tokyo), 68: 763–6.

Shrestha, A., Sultana, R, Chae, J.C., Kim K., and Lee, K.J. 2015. ‘Bacillus thuringiensis C25 which is rich in cell wall degrading enzymes efficiently controls lettuce drop caused by Sclerotinia minor’, European Journal of Plant Pathology, 142: 577–89.

Silby, M. W., A. M. Cerdeno-Tarraga, G. S. Vernikos, S. R. Giddens, R. W. Jackson, G. M. Preston, X. X. Zhang, C. D. Moon, S. M. Gehrig, S. A. Godfrey, C. G. Knight, J. G. Malone, Z. Robinson, A. J. Spiers, S. Harris, G. L. Challis, A. M. Yaxley, D. Harris, K. Seeger, L. Murphy, S. Rutter, R. Squares, M. A. Quail, E. Saunders, K. Mavromatis, T. S. Brettin, S. D. Bentley, J. Hothersall, E. Stephens, C. M. Thomas, J. Parkhill, S. B. Levy, P. B. Rainey, and N. R. Thomson. 2009. ‘Genomic and genetic analyses of diversity and plant interactions of Pseudomonas fluorescens’, Genome Biol, 10: R51.

Taghavi, S., D. van der Lelie, A. Hoffman, Y. B. Zhang, M. D. Walla, J. Vangronsveld, L. Newman, and S. Monchy. 2010. ‘Genome sequence of the plant growth promoting endophytic bacterium Enterobacter sp. 638’, PLoS Genet, 6: e1000943.

Tapia, J., R. Gonzalez, B. Townley, V. Oliveros, F. Alvarez, G. Aguilar, A. Menzies, and M. Calderon. 2018. ‘Geology and geochemistry of the Atacama Desert’, Antonie Van Leeuwenhoek, 111: 1273–91.

Undabarrena, A., F. Beltrametti, F. P. Claverias, M. Gonzalez, E. R. Moore, M. Seeger, and B. Camara. 2016. ‘Exploring the Diversity and Antimicrobial Potential of Marine Actinobacteria from the Comau Fjord in Northern Patagonia, Chile’, Front Microbiol, 7: 1135.

Uritskiy, G., and J. DiRuggiero. 2019. ‘Applying Genome-Resolved Metagenomics to Deconvolute the Halophilic Microbiome’, Genes (Basel), 10.

Uritskiy, G., S. Getsin, A. Munn, B. Gomez-Silva, A. Davila, B. Glass, J. Taylor, and J. DiRuggiero. 2019. ‘Halophilic microbial community compositional shift after a rare rainfall in the Atacama Desert’, ISME J, 13: 2737–49.

Valdes Franco, J. A., R. Collier, Y. Wang, N. Huo, Y. Gu, R. Thilmony, and J. G. Thomson. 2016. ‘Draft Genome Sequence of Agrobacterium rhizogenes Strain NCPPB2659’, Genome Announc, 4.

Vlamakis, H., Y. Chai, P. Beauregard, R. Losick, and R. Kolter. 2013. ‘Sticking together: building a biofilm the Bacillus subtilis way’, Nat Rev Microbiol, 11: 157–68.

Wang, X. Q., C. M. Li, C. A. Dunlap, A. P. Rooney, and Z. J. Du. 2018. ‘Marinicella sediminis sp. nov., isolated from marine sediment’, Int J Syst Evol Microbiol, 68: 2335–39.

Weon, H. Y., S. W. Kwon, J. A. Son, S. J. Kim, Y. S. Kim, B. Y. Kim, and J. O. Ka. 2010. ‘Adhaeribacter aerophilus sp. nov., Adhaeribacter aerolatus sp. nov. and Segetibacter aerophilus sp. nov., isolated from air samples’, Int J Syst Evol Microbiol, 60: 2424–9.

Xiaomei Y, Yao T. 2020. ‘Comparative genomic analysis of Bacullus mycoides Gnyt1 strain’, Epigenetic and Genomics, preprint.

Yoon, S. H., S. M. Ha, S. Kwon, J. Lim, Y. Kim, H. Seo, and J. Chun. 2017. ‘Introducing EzBioCloud: a taxonomically united database of 16S rRNA gene sequences and whole-genome assemblies’, Int J Syst Evol Microbiol, 67: 1613–17.

Zachow, C., H. Muller, J. Monk, and G. Berg. 2017. ‘Complete genome sequence of Pseudomonas brassicacearum strain L13-6-12, a biological control agent from the rhizosphere of potato’, Stand Genomic Sci, 12: 6.

Zhao, Y., J. N. Selvaraj, F. Xing, L. Zhou, Y. Wang, H. Song, X. Tan, L. Sun, L. Sangare, Y. M. Folly, and Y. Liu. 2014. ‘Antagonistic action of Bacillus subtilis strain SG6 on Fusarium graminearum’, PLoS One, 9: e92486.

